# Remote control of neural function by X-ray-induced scintillation

**DOI:** 10.1101/798702

**Authors:** Takanori Matsubara, Takayuki Yanagida, Noriaki Kawaguchi, Takashi Nakano, Junichiro Yoshimoto, Maiko Sezaki, Hitoshi Takizawa, Satoshi P. Tsunoda, Shin-ichiro Horigane, Shuhei Ueda, Sayaka Takemoto-Kimura, Hideki Kandori, Akihiro Yamanaka, Takayuki Yamashita

## Abstract

Scintillators emit visible luminescence when irradiated with X-rays. Given the unlimited tissue penetration of X-rays, the employment of scintillators could enable remote optogenetic control of neural functions at any depth of the brain. Here we show that a yellow-emitting inorganic scintillator, Ce-doped Gd3(Al,Ga)5O12 (Ce:GAGG), could effectively activate red-shifted excitatory and inhibitory opsins, ChRmine and GtACR1, respectively. Using injectable Ce:GAGG microparticles, we successfully activated and inhibited midbrain dopamine neurons in freely moving mice by X-ray irradiation, producing bidirectional modulation of place preference behavior. Ce:GAGG microparticles were non-cytotoxic and biocompatible, allowing for chronic implantation. Pulsed X-ray irradiation at a clinical dose level was sufficient to elicit behavioral changes without reducing the number of radiosensitive cells in the brain and bone marrow. Thus, scintillator-mediated optogenetics enables less invasive, wireless control of cellular functions at any tissue depth in living animals, expanding X-ray applications to functional studies of biology and medicine.

---

Optogenetics has enabled the elucidation of the causal roles of specific neurons in driving circuit dynamics, plasticity, and behavior^1,2^. Clinical treatment of neurological diseases would also benefit from optogenetic approaches that can control functions of well-defined neural circuits in precise timings^3-7^. However, the application of optogenetics to thick and large tissues such as human brains has been hampered because the stimulating light (wavelength: ∼430–610 nm) used to activate light-sensitive proteins is heavily scattered and absorbed by tissues^1^. Optogenetic stimulation of neurons deep in the brain usually requires the invasive implantation of optical fibers tethered to an external light source. These tethered fiber optics, although widely employed, are known to pose diverse problems, including tissue damage, neuroinflammatory responses, phototoxicity and thermal effects upon irradiation, as well as physical restriction of animal movement^8-11^. Recent studies have shown that injectable upconverting nano/microparticles, which emit visible light in response to tissue-penetrating near-infrared (NIR) light irradiation, can be used for minimally invasive actuation of neurons deep in the brain^12-16^. However, even NIR light penetrates only up to several millimeters of tissue. Furthermore, the low upconversion yields of these particles demand high-energy NIR illumination which can cause abrupt tissue heating and photodamage^15,16^. Considering the fact that therapeutic targets for human deep brain stimulation can be more than several centimeters deep, non-optical forms of energy delivery should be pursued. In fact, methods to control the activities of specific neuronal populations using magneto-thermal^17^ and ultrasonic^18^ stimulation have been explored; however, these approaches are associated with a significantly reduced time resolution compared with optogenetics and are currently restricted by limited compatibility with free behavior.

Here, we report the development of an X-ray-mediated, wireless optogenetic technology, which is practically unconstrained by tissue depth. We employ a scintillator that can absorb the energy of incoming X-ray particles and release it in the form of visible luminescence called scintillation. Scintillators are widely used in various particle detectors, such as X-ray security and computed tomography (CT) scanners. However, it has not yet been proven whether scintillation can effectively actuate neurons in living animals by activating light-sensitive proteins. Also, it is currently unknown whether scintillators are biocompatible and can be safely implanted in living animals. We found that an inorganic scintillator, Ce-doped Gd3(Al,Ga)5O12 (Ce:GAGG), emits yellow scintillation^19,20^ to effectively activate red-shifted excitatory and inhibitory opsins. Using Ce:GAGG microparticles, we were able to actuate a specific neuronal population in freely moving mice, driving related behaviors with X-ray irradiation that does not harm radiosensitive cells. Our analysis also shows the biocompatibility of Ce:GAGG microparticles. Overall, this work demonstrates that X-rays can be used to control the function of cells at any tissue depth, expanding the range of X-ray applications in biology and medicine.

## Results

### Ce:GAGG luminescence activates red-shifted opsins

When irradiated onto a mouse head (Fig. 1a and Supplementary Fig. 1), X-rays readily penetrated through the head skin, skull, and brain tissue, whereas most of the energy derived from NIR and visible light did not reach deep in the brain due to absorption and scattering by tissue. Moreover, X-ray irradiation did not increase the temperature of tissues, whereas NIR illumination with a conventional optogenetic stimulation protocol caused striking tissue heating (Supplementary Fig. 2). These results highlight the distinct advantages of using X-ray-induced scintillation for remote optogenetic control of neural circuits deep in the brain. In this study, we utilized single scintillator crystals of Ce:GAGG which emit yellow luminescence in response to UV or X-ray radiation^19,20^ (Fig. 1b). These crystals were transparent and non-deliquescent (Fig. 1b). UV-induced photo-luminescence (PL) and X-ray-induced radio-luminescence (RL) of a Ce:GAGG crystal have essentially the same spectrum^19,20^ (peak wavelength: 520–530 nm; Fig. 1b) because both PL and RL are based on the 5d-4f transitions of Ce3+. The RL light yield of Ce:GAGG is reported to be 46,000 photons/MeV^19,20^.

**Fig. 1.**
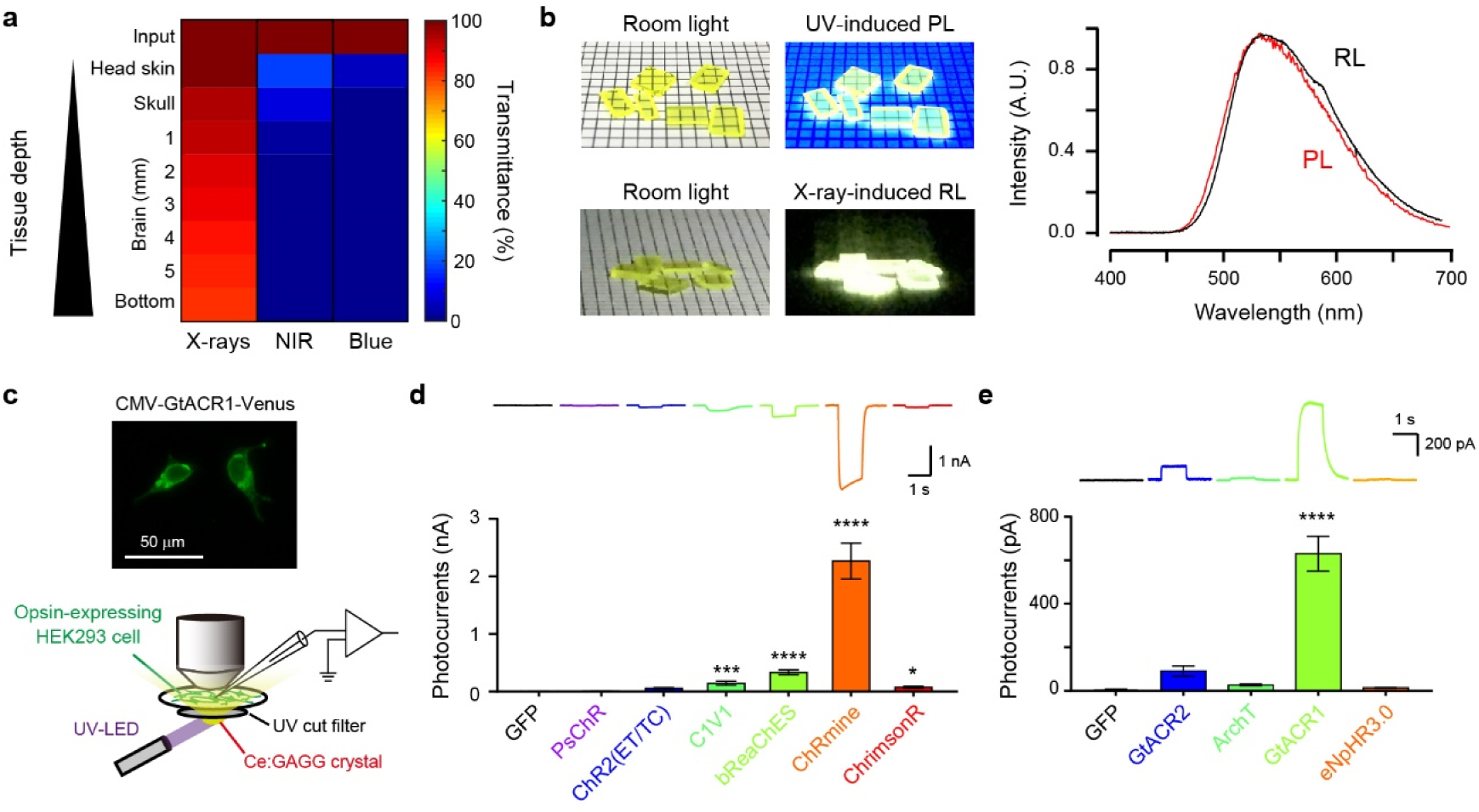
Potential of Ce:GAGG scintillation for application in deep brain optogenetics. **a**, Tissue transmittance of X-rays (150 kV, 3 mA), NIR (976 nm), and blue light (470 nm) irradiated onto a mouse head. Values were estimated using the data shown in Supplementary Fig. 1. **b**, Left, Ce:GAGG crystals under room light. Middle, Ce:GAGG crystals under UV (365 nm) and X-ray (150 kV, 3 mA). Right, spectra of RL (black) and PL (red) emitted by a single Ce:GAGG crystal. **c**, Top, epi-fluorescence image of HEK 293 cells expressing GtACR1-Venus. Bottom, schematic of the photo-current recording. **d**, Representative photocurrents (top) and photocurrent amplitudes (bottom) of different excitatory opsins and GFP induced by Ce:GAGG PL (*n* = 10 to 14 cells, * * * * *P* < 0.0001, * * * *P* < 0.001, * *P* < 0.05, Dunn’s multiple comparison tests vs. GFP). PL power: 1.8 mW/cm^2^. **e**, Same as (d) but with inhibitory opsins and GFP (*n* = 10 to 12 cells, Dunnett’s multiple comparison tests vs. GFP, * * * * *P* < 0.0001). PL power: 1.8 mW/cm^2^. Values are mean ± SEM.

We first sought opsins that could be effectively activated by the PL of Ce:GAGG. DNA plasmids encoding different opsins were transfected into HEK 293 cells and photocurrents induced by PL illumination were measured (Fig. 1c). For depolarizing opsins, the yellow PL of Ce:GAGG (1.8 mW/cm^2^) elicited the largest photocurrents in cells expressing the red-shifted opsin ChRmine^21^ (2.28 ± 0.31 nA, *n* = 11; Fig. 1d). Regarding inhibitory opsins, the anion channelrhodopsin GtACR1^22^ showed the strongest activation (627.2 ± 80.2 pA, *n* = 12; Fig. 1e) in response to the PL of Ce:GAGG. PL-irradiation of GFP-expressing cells induced undetectable currents (Fig. 1d,e).

Thus, Ce:GAGG PL can activate red-shifted opsins that are used for optogenetic control of neurons. Furthermore, the yellow PL of Ce:GAGG could also induce significant activation of the enzyme rhodopsin BeGC1^23^ (Supplementary Fig. 3), suggesting that the scintillation of Ce:GAGG could be used for the activation of various light-sensitive proteins.

### Bidirectional control of neuronal activities by Ce:GAGG luminescence

We next examined the intensity of Ce:GAGG luminescence required to actuate neuronal activities. Cre-dependent adeno-associated virus (AAV) vectors were injected into the ventral tegmental area (VTA) of DAT-IRES-Cre mice to induce the specific expression of ChRmine in dopamine (DA) neurons (Fig. 2a and Supplementary Fig. 4). In acute slice preparations, 1-s pulses of the yellow PL of Ce:GAGG elicited depolarizing photocurrents in DA neurons in a PL intensity-dependent manner (Fig. 2b). Irradiation with 3.3 μW/cm^2^ elicited action potentials (APs) in 9 out of 15 DA neurons, and the rate of PL-evoked APs plateaued at approximately 15 μW/cm^2^ (Fig. 2c,d). When the cells were held at around - 40 mV to exhibit spontaneous APs, 1-min PL illumination (Fig. 2e,f), and 1-s pulsed PL illumination (5–20 Hz; Supplementary Fig. 5) increased the AP rate at an even lower intensity (1.7 μW/cm2). In DA neurons expressing inhibitory soma-targeted GtACR1^24^ (stGtACR1; Supplementary Fig. 4), hyperpolarizing photocurrents were induced by Ce:GAGG PL illumination (Fig. 2g), which attenuated AP generation at 1.7 μW/cm^2^ and maximally suppressed spiking at around 15 μW/cm^2^ PL (Fig. 2h–k). Thus, the activity of ChRmine- or stGtACR1-expressing neurons can be modulated by illumination with Ce:GAGG PL at a few microwatts intensities.

**Fig. 2.**
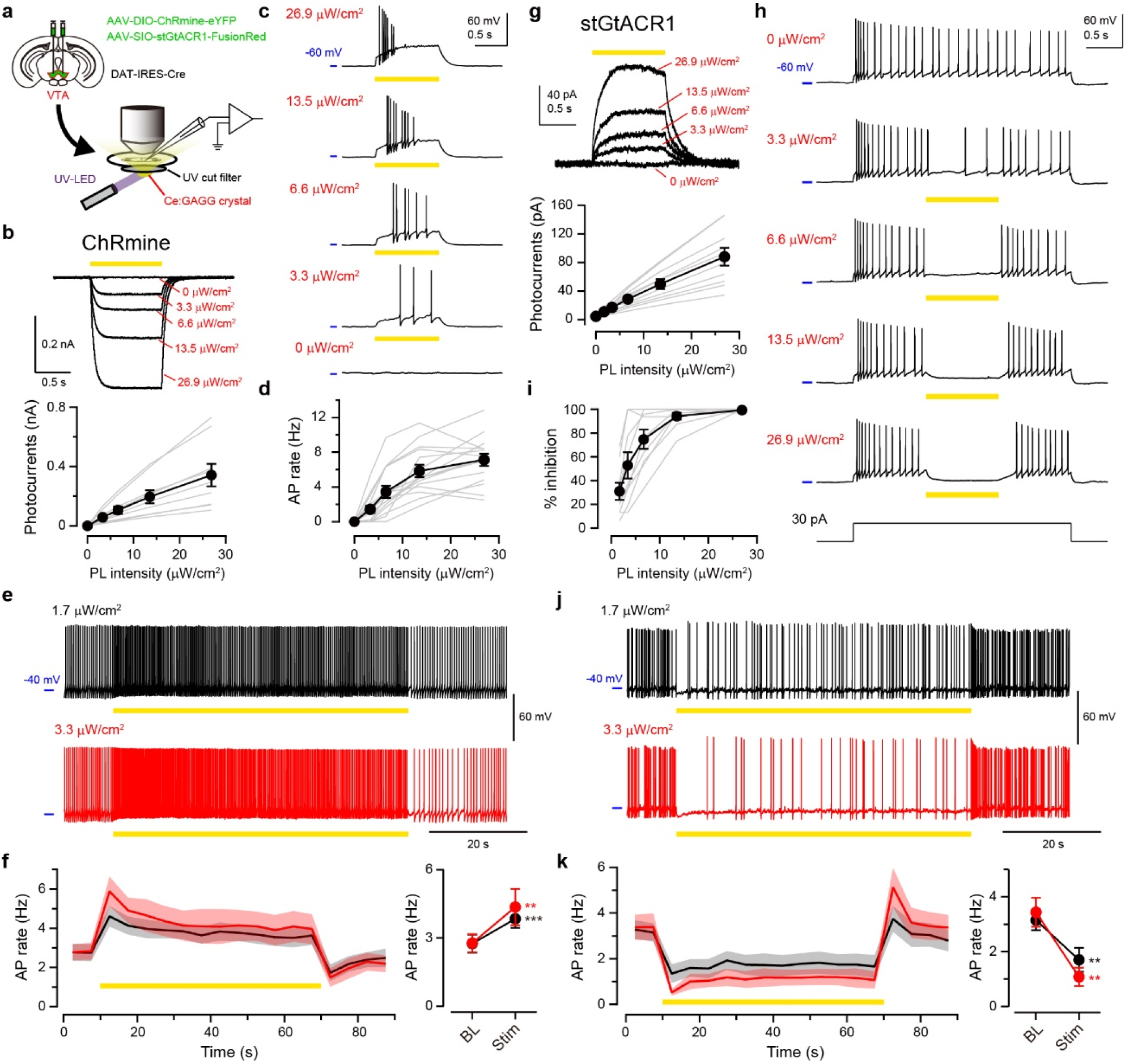
Photo-luminescence of Ce:GAGG bidirectionally actuated VTA-DA neurons *in vitro*. **a**, Schematic of the experiment. **b**, Top, sample voltage-clamp recordings from a ChRmine-expressing VTA-DA neuron responding to 1-s Ce:GAGG PL. Bottom, photocurrent amplitude vs. PL intensity (*n* = 9 cells). **c**, Sample recordings from a ChRmine-expressing VTA-DA neuron current-clamped at approximately −60 mV, responding to 1-s PL. **d**, AP rate of ChRmine-expressing neurons vs. PL intensity (*n* = 15 cells). **e**, Sample recordings from a ChRmine-expressing neuron current-clamped at approximately −40 mV, responding to 1-min PL illumination. **f**, Time course (left) and quantification (right) of average AP rates (*n* = 7 cells) with 1-min PL illumination at 1.7 (black) or 3.3 (red) μW/cm^2^. The shadows indicate SEM. BL: baseline, Stim: stimulation. **g**, Same as **b**, but from stGtACR1-expressing VTA-DA neurons (*n* = 11 cells). **h**, Same as **c**, but for a stGtACR1-expressing neuron. APs were evoked by current injection. **i**, Success rate of spike suppression by Ce:GAGG PL in stGtACR1-expressing neurons (*n* = 10 cells). **j, k**, Same as **e, f** but for stGtACR1-expressing neurons (*n* = 7 cells). Gray lines indicate individual cells. * * * *P* < 0.001, * * *P* < 0.01, paired *t*-test. Values are mean ± SEM.

### Ce:GAGG microparticles enable neuronal actuation *in vivo*

Having identified scintillator-opsin combinations that modulate neuronal activity, we next examined the ability of X-ray-induced RL of the Ce:GAGG crystal to activate neurons in vivo. We first pulverized the Ce:GAGG crystal into particles with an average size of 2.3 μm (named scintillator microparticles, SMPs; Fig. 3a), to be injected into the brain. The intensity of RL emitted by SMPs, which was proportional to the dose rate of X-irradiation (Supplementary Fig. 6), can reach ∼1.5–3 μW/cm^2^ near the injection site in vivo with X-irradiation at a rate of 1.0 Gy/min (Fig. 3b). We next induced the expression of ChRmine in VTA-DA neurons using AAV injections and injected the SMPs (50 mg/ml, 600 nl) at the same location as the AAV injection (Fig. 3c). X-irradiation for a total of 5 min (1-min pulses every 2 min, five times, at 0.5 or 1.0 Gy/min) induced c-Fos expression in a larger fraction of ChRmine-expressing neurons compared to control conditions (Fig. 3d,e and Supplementary Fig. 7). The fraction of c-Fos-positive cells increased as the rate of X-irradiation increased. Thus, the SMPs injected in vivo can activate neurons in an X-ray- and opsin-dependent manner.

**Fig. 3.**
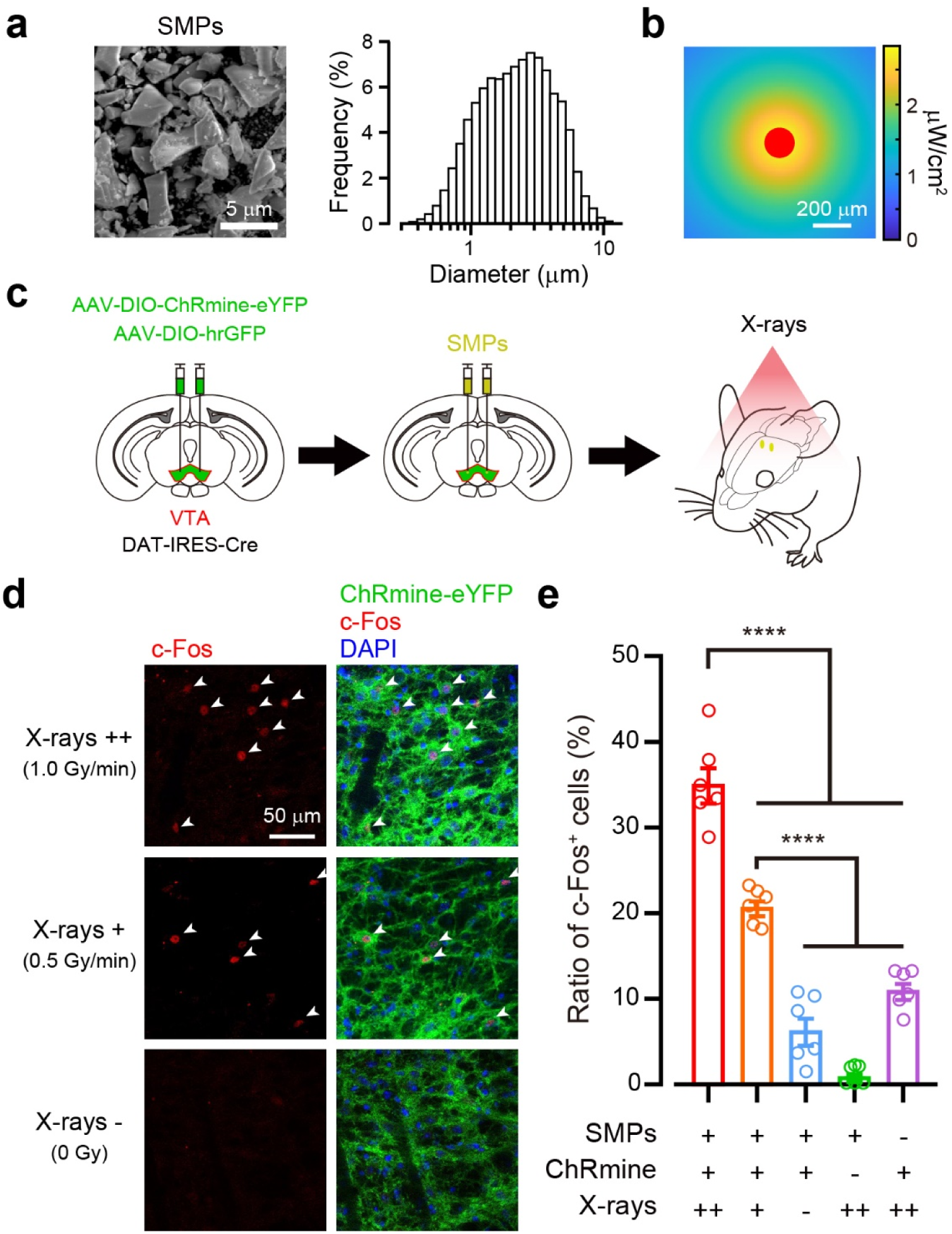
Radio-luminescence of Ce:GAGG microparticles activated VTA-DA neurons *in vivo*. **a**, Left, a scanning electron micrograph of SMPs. Right, size distribution of the SMPs measured by dynamic light scattering. **b**, An RL intensity map around the SMPs (red) injected in gray matter under 1.0 Gy/min X-irradiation (see also Supplementary Fig. 6). **c**, Schematic of the c-Fos induction experiment. **d**, Representative confocal images showing RL-induced expression of c-Fos (red, arrow heads) among ChRmine-expressing cells near injected SMPs. **e**, Ratio of c-Fos-positive cells under different conditions (*n* = 6 hemispheres from 3 mice for each group). * * * * *P* < 0.0001, Bonferroni’s multiple comparison test. Open circles indicate individual data. Values are mean ± SEM.

We further assessed the biocompatibility of the scintillator. The vast majority of dissociated hippocampal neurons around the Ce:GAGG crystal in the culture dish survived for 7 days (Fig. 4a), and a fraction of neurites extended towards the crystal (Fig. 4a). Similarly, HEK 293 cells cultured in the presence of the Ce:GAGG crystal proliferated at a normal rate (Supplementary Fig. 8). Furthermore, SMPs implanted into the brain for one week caused less activation of microglial cells than that caused by implantation of an optical fiber (Supplementary Fig. 9). Notably, the number of neurons was significantly reduced around the implanted optic fibers (one week after surgery, 11.9% of control, *P* = 0.0005; four weeks after surgery, 25.5% of control, *P* = 0.003; Fig. 4b,c), whereas that was unchanged around injected SMPs (one week after surgery, 88.1% of control, *P* = 0.90; four weeks after surgery, 100% of control, *P* = 1.0; Fig. 4b,c). Thus, the Ce:GAGG crystal is biocompatible and can be implanted for a long period without cytotoxicity.

**Fig. 4.**
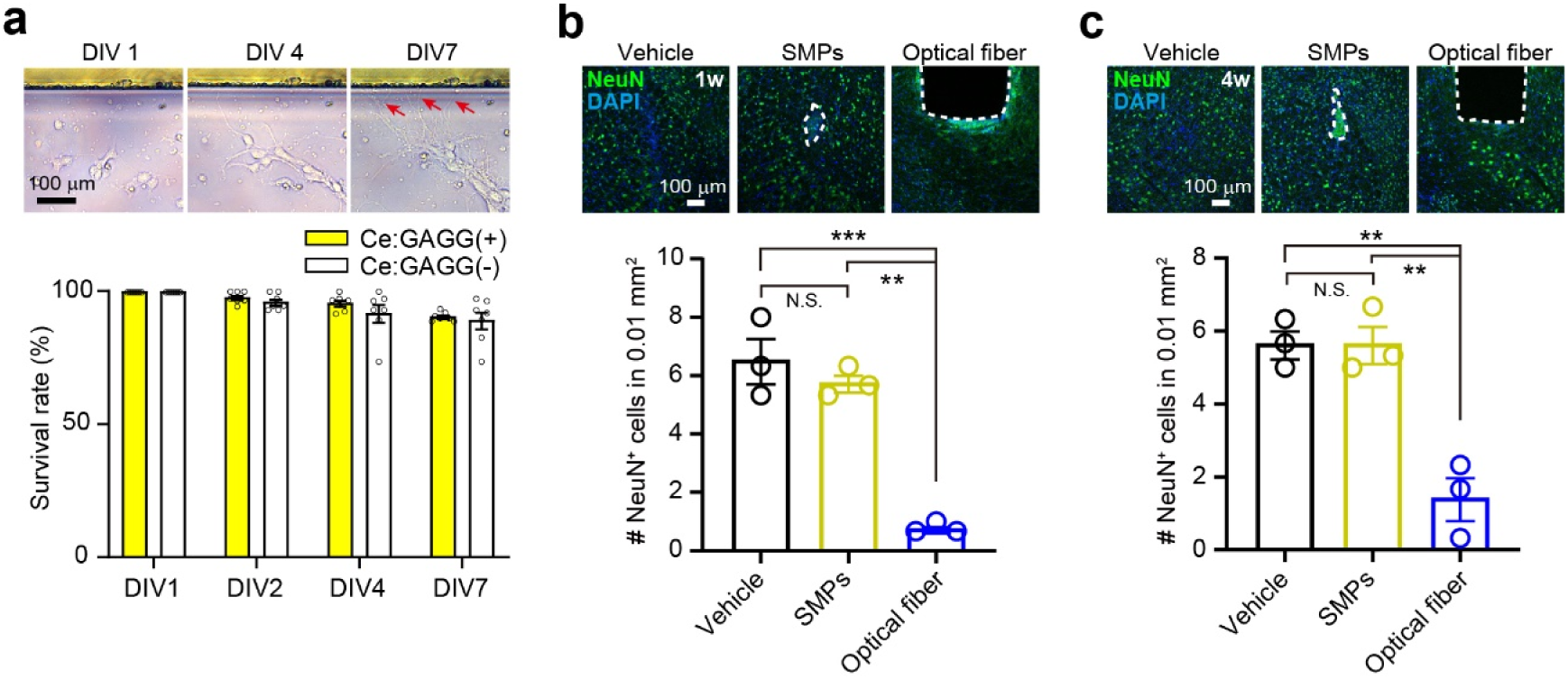
Ce:GAGG microparticles were non-cytotoxic and biocompatible. **a**, Top, dissociated hippocampal neurons cultured with a Ce:GAGG crystal (upper). Neurites extend towards the crystal (arrows) at 7 days *in vitro* (DIV). Bottom, the survival rates of dissociated neurons cultured with or without a Ce:GAGG crystal (*n* = 7 dishes for each group, *P* > 0.15). **b, c**, Top, representative confocal images of immunoreactivity against NeuN (green) at the injection site of vehicle (left) or SMPs (middle), and at the ventral tip of an implanted optical fiber (right) at one week (**b**) or four weeks (**c**) after surgery. The trace of SMPs or an optical fiber is outlined by a dashed line. Blue: DAPI. Bottom, average number of NeuN-positive cells counted in a 100 μm x 100 μm square near the injection/implantation traces (*n* = 3 mice for each group). Neuronal loss was found near the optical fiber traces, but not at around implanted SMPs. * * * *P* < 0.001, * * *P* < 0.01, * * *P* < 0.01, Bonferroni’s multiple comparison test. N.S., not significant. Open circles indicate individual data.Values are mean ± SEM.

### Bidirectional change of behaviors induced by scintillator-mediated optogenetics

We finally tested whether the scintillator-mediated actuation of neurons in vivo could induce behavioral changes. Transient activation and inhibition of DA neurons in the VTA are sufficient for behavioral conditioning^25-27^. We therefore induced the expression of excitatory ChRmine or inhibitory stGtACR1 in VTA-DA neurons through viral injections, and bilaterally injected SMPs in the VTA (Fig. 5a). The conditioned place preference (CPP) test was performed by placing the mice into a test chamber with two compartments, only one of which was irradiated with either X-ray pulses (50 ms, 10 Hz, 10 times every 30 s) through an X-ray chopper wheel (“Pulsed” conditioning [P.C.]; Fig. 5b,c and Supplementary Fig. 10) or continuous X-ray irradiation (“Free moving” conditioning [F.C.]; Fig. 5d, and Supplementary Fig. 10). The initial place preference was not different between the opsin-expressing and GFP-expressing control mice. After P.C., however, mice expressing ChRmine had a significantly higher preference for the X-ray conditioned compartment than control mice (Fig. 5d,e), whereas those expressing stGtACR1 had a lower preference for the conditioned compartment after F.C. (Fig. 5f–i). Thus, scintillator-mediated remote optogenetics can be used for bidirectional neuronal actuation deep in the brain of mice, resulting in behavioral changes.

**Fig. 5.**
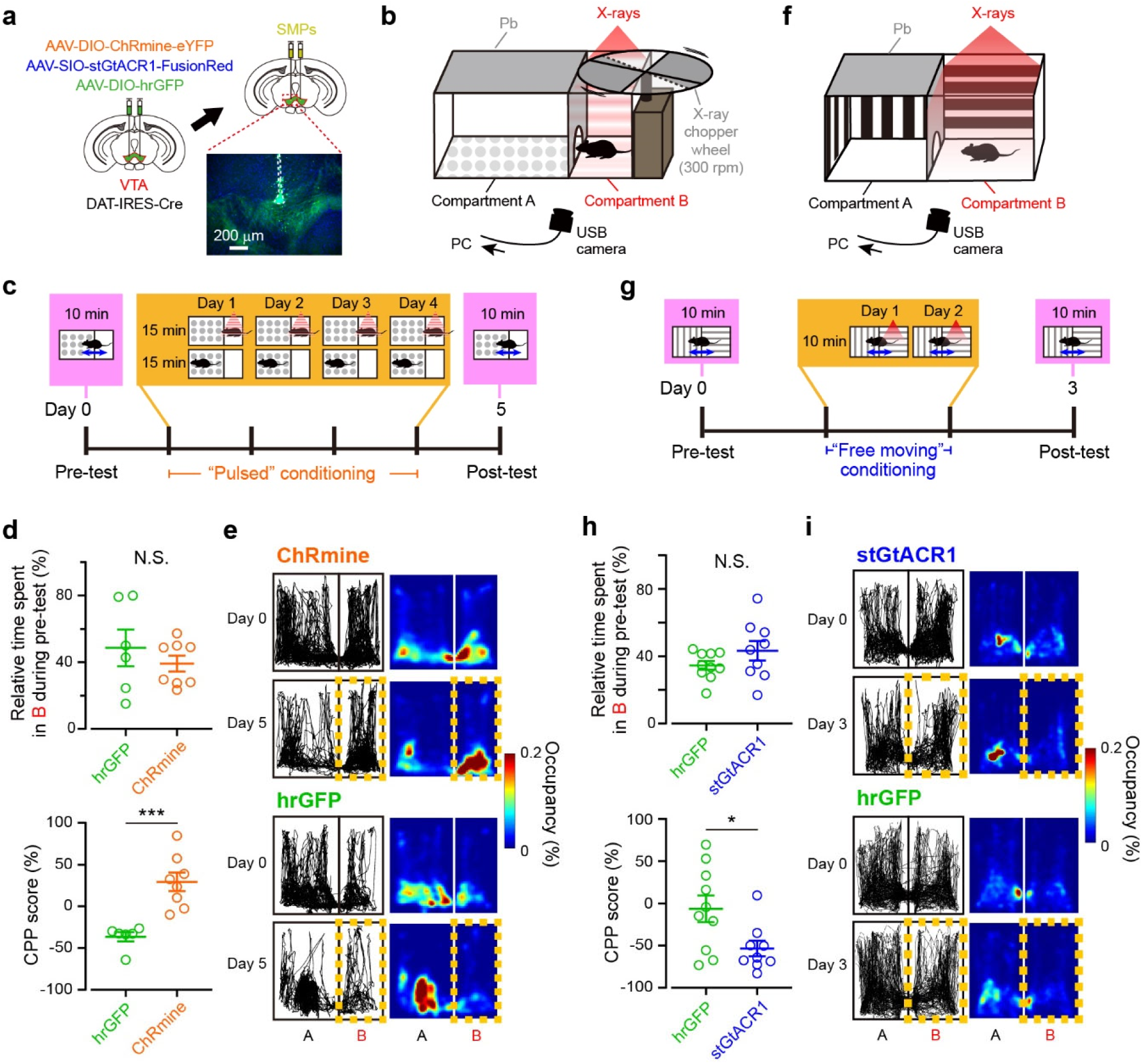
Scintillator-mediated wireless optogenetics drove conditioned place preference and aversion in freely behaving mice. **a**, Schematic of the experiment. Inset: an epi-fluorescence image showing SMPs (dashed outline) injected at dorsal VTA. Green: ChRmine-eYFP, Blue: DAPI. **b**, CPP test chamber for “Pulsed” conditioning. The X-ray shieldings and the X-ray wheel chopper are made of lead (Pb). **c**, Time course of CPP test with “Pulsed” conditioning. The gate between two compartments is open for pre- and post-tests but closed during “Pulsed” conditioning. **d**, Quantification of CPP with “Pulsed” conditioning (*n* = 6 mice for hGFP group, 8 mice for ChRmine group). **e**, Representative tracking data (left) and the corresponding heat maps (right) for mice before (Day 0) and after (Day 5) “Pulsed” conditioning, as extracted from oblique-view movies. **f**, CPP test chamber for “Free moving” conditioning. **g**, Time course of CPP test with “Free moving” conditioning. The gate between two compartments is open throughout the test. **h**, Quantification of CPP with “Free moving” conditioning (*n* = 10 mice for hGFP group, 9 mice for stGtACR1 group). **i**, Same as **e**, but for mice before (Day 0) and after (Day 3) “Free moving” conditioning. * * * *P* < 0.001, * *P* < 0.05, unpaired *t* test. N.S., not significant. Open circles indicate individual data. Values are mean ± SEM.

The total dose of X-irradiation during the behavioral tests is far below the threshold for acute neuronal and vascular dysfunction in the brain^28,29^. In fact, X-irradiation corresponding to the F.C. (∼7 Gy) did not change locomotor behavior (Supplementary Fig. 11) and blood brain barrier function (Supplementary Fig. 12). In a long-term observation, all the mice that experienced the F.C. survived for at least 8 weeks after radiation with no significant difference in body weight from non-X-irradiated control mice at the same age (Supplementary Fig. 13a). These results suggest that scintillator-mediated optogenetics can be used for days-long behavioral experiments in mice even with a relatively high dose of radiation. On the other hand, the total body X-irradiation is known to damage radiosensitive cell populations, depending on its cumulative dose. A high-dose X-irradiation reduced the number of immature neurons in the hippocampal dentate gyrus (Supplementary Fig. 14) through apoptotic cell death of neuronal precursor cells^30,31^ (Supplementary Fig. 15). In consistent with previous reports^30,31^, hippocampal neurogenesis was partially impaired at 8 weeks after the F.C. radiation (Supplementary Fig. 13b–f). However, the P.C., which causes considerably less radiation dose (∼0.5 Gy), induced neither loss of immature neurons (Supplementary Fig. 14) nor apoptosis (Supplementary Fig. 15) in the hippocampus. Although a high-dose X-irradiation caused an overall reduction in the number of bone marrow cells^32^ without specificity of cell-types (Supplementary Fig. 16), the P.C. rather increased the number of the total bone marrow cells (Supplementary Fig. 16) presumably due to radio-resistance effects^33^. Thus, reducing radiation dose using pulsed X-irradiation extends the utility of scintillator-mediated optogenetics to investigation over a larger timescale.

## Discussion

We have here demonstrated the feasibility of a novel, scintillator-mediated optogenetic technology that allows full wireless control of neuronal activity in behaving animals. The scintillator particles were biocompatible and injectable, and remained at the injection site for long periods without cytotoxicity, serving as minimally invasive optogenetic actuators. The thermal effects on neuronal activity were negligible using this technology (Supplementary Fig. 2), a significant advantage over conventional^9,11^ and NIR-mediated^15,16^ optogenetics. Using a careful radiation dose setting, this technology can be safely applied in a variety of rodent behavioral experiments, which are normally hindered by the tethered fiber optics, or the large implant on the head in other wireless optogenetic technologies^8,34^.

Given the unlimited tissue penetration of X-rays, scintillator-mediated optogenetics can be readily applied to larger animals, including monkeys. Most behavioral experiments combined with electrophysiology in monkeys are performed under head-restrained conditions. In such experiments, focal X-irradiation of the brain, which prevents radiation exposure to other organs, would be feasible. The total radiation dose of our P.C. procedure (∼0.5 Gy), which is roughly equivalent to the dose of a single perfusion CT scan^35,36^, is more than 100 times lower than the standard dose of radiotherapy for brain tumor treatment (50–70 Gy^28,29^). Although we have shown that scintillator-mediated optogenetics can be used for remote actuation of neurons using X-irradiation at a non-toxic cumulative dose, increased safety in the use of this technology can be achieved by stereotactic focusing of small radiation beams on specific brain regions, as was achieved in gamma knife surgery^37^.

Since Röntgen’s discovery in the late 19th century^38^, X-rays have been widely used for medical imaging and cancer therapy. However, X-rays have never been used to control the physiological functions of cells in living animals, as we have shown here. The development of scintillator-mediated optogenetics thus expands the application of X-rays to functional studies of biology and medicine and opens new avenues for therapeutic strategies using optogenetics. Biomedical technologies that use visible light for genome editing^39^ or control of intracellular signaling^40^ would benefit from wireless applications targeting deeper tissues, which is now possible with scintillator-mediated approaches. Future improvements in light yields of scintillators, engineering of opsin-bound scintillator nanocrystals, and combination with focused X-irradiation will all contribute to allowing control of cellular functions over larger volumes of tissue with less risk of radiation toxicity.

## Methods

### Scintillator preparation

The single crystal Ce:GAGG was synthesized using the conventional Czochralski method in Furukawa^19,20^. For electrophysiology (Fig. 1 and 2, and Supplementary Fig. 5) and other measurements (Supplementary Fig. 2, 3 and 8), the crystal was fabricated into 3−8 mm rectangular blocks 0.5−1 mm thick. For injection of the crystals in the mouse brain, we pulverized the Ce:GAGG crystals into particles using a planetary ball mill. These particles were further crushed in an agate mortar and collected in ethanol. The ethanol solution containing the particles was sonicated for 10 min, and smaller particles were obtained from the supernatant after removal of precipitate. The solution was then centrifuged at 13,000 rpm for 30 s, and supernatant was discarded. Ethanol was added to the precipitate containing Ce:GAGG particles. We repeated the sonication of the solution, the removal of precipitate and the centrifugation to remove supernatant for total three times. The final precipitate contained micro-sized Ce:GAGG particles (SMPs). After evaporation of the remaining vehicle, SMPs (50 mg/ml) were dispersed with Ringer’s solution containing (in mM) 135 NaCl, 5 KCl, 5 HEPES, 1.8 CaCl_2_, 1 MgCl_2_ (adjusted to pH 7.3 with NaOH).

### Luminescence measurements

The PL of a Ce:GAGG crystal was induced with 365-nm UV light (LEDMOD V2, Omicron; LC-L2, Hamamatsu) unless otherwise noted. The PL intensity for electrophysiological recordings was routinely measured with a photodiode sensor (1 cm × 1 cm; PD300-1W, Ophir) placed over the recording chamber through a UV-cut filter fabricated from UV-cut goggles (SSUV 297, AS ONE) on each experimental day. In experiments using acute slices, the PL intensity over the slice in the recording chamber was measured. The intensity of unfiltered UV light over the recording chamber was <0.1 μW/cm^2^. The RL power of SMPs (Supplementary Fig.6) was measured with a fiber-coupled photoreceiver (Newport 2151) through an optical fiber (tip diameter: 400

μm) placed close to a mass of SMPs exposed to X-ray irradiation in an X-ray machine (MX-160Labo, mediXtec). The photoreceiver current, which was sampled using an analogue-digital converter (Picoscope 4262, Pico Technology), was calibrated against the PL power measured by the photodiode sensor and converted to luminescence intensity (in watts). For 3D simulation of the RL intensity (Fig. 3b), we assumed that the average RL intensity at the surface of a spherical aggregate of the SMPs corresponds to the measured RL power (Supplementary Fig.6). PL emission spectra under UV illumination (340 ± 10 nm, LAX-103, Asahi Spectra) were measured using a spectrometer (QE-Pro, Ocean Optics). RL emission spectra under X-ray irradiation (70 kV, 1 mA) were measured through an optical fiber using a CCD spectrometer (DU-420-BU2, Andor)^20^.

### Animals

All experiments were performed in accordance with the guidelines of the Physiological Society of Japan and approved by the institutional review board of the Research Institute of Environmental Medicine, Nagoya University, Japan. Adult C57BL6/J mice and DAT-IRES-Cre mice (B6.SJL-Slc6a3^tm1.1(cre)Bkmn^/J, The Jackson Laboratory) of both sexes were maintained on a 12-h light-dark cycle, with free access to food and water. DAT-IRES-Cre mice were maintained as homogenic mutants. Only 11−18-week-old male DAT-IRES-Cre mice were used for the CPP experiments.

### Plasmids

For expression in HEK293 cells, all plasmids encoding opsins and fluorescent proteins were constructed by subcloning into an empty pCMV vector unless otherwise noted. pCMV-PsChR-Venus and pCMV-C1V1-Venus were obtained from H. Yawo (Tohoku University). *bReaChES-TS-eYFP* and *ChRmine-eYFP* were isolated from pAAV-CaMKIIa-DIO-bReaChes-TS-eYFP and pAAV-CaMKIIa-DIO-ChRmine-eYFP, respectively, both of which were gifted by K. Deisseroth (Stanford University). A full-length gene encoding BeGC1 (accession number KF309499) was synthesized after human codon optimization and inserted into the peGFP-N1 vector. For AAV production, pAAV-Ef1a-DIO-ChRmine-eYFP-WPRE was obtained from K. Deisseroth (Stanford University) and pAAV-hSyn1-SIO-stGtACR1-FusionRed was obtained from Addgene (#105678).

### Cell culture and transfection

For electrophysiological recordings in cultured cells, expression vector plasmids encoding opsins or hrGFP were transfected into HEK 293 cells using Lipofectamine 2000 (Thermo Fisher Scientific). The cells were washed with phosphate-buffered saline (PBS) 3–4 hours after transfection, and then seeded on a coverslip (12 mm diameter) in Dulbecco’s modified Eagle’s medium (DMEM; Sigma-Aldrich) supplemented with 10% (vol/vol) fetal bovine serum (FBS), 100 U/ml penicillin, and 0.1 mg/ml streptomycin. The cells were maintained in the medium in an incubator at 37°C and 5% CO_2_/95% air for 24−36 hours before recordings.

Dissociated hippocampal neurons were prepared from embryonic (E17.5) mice. Isolated hippocampal tissues were incubated with Hank’s Balanced Salt Solution (HBSS; Sigma-Aldrich) containing 1% DNase I (Sigma-Aldrich) and 2.5% trypsin for 10 min at 37°C, and then washed three times with HBSS. The tissues were then dispersed by pipetting in Neurobasal medium (Thermo Fisher Scientific). After removing aggregated cells by filtration, the hippocampal cells were seeded on a coverslip (12 mm diameter) coated with poly-L-lysine in DMEM (Sigma-Aldrich) and incubated for 4 h at 37°C. The culture medium was subsequently replaced by Neurobasal medium supplemented with 0.5 mM GlutaMAX (Thermo Fisher Scientific), 2% (vol/vol) B-27 (Thermo Fisher Scientific), 100 U/ml penicillin and 0.1 mg/ml streptomycin. The cultured neurons were maintained in the medium in an incubator at 34°C with 5% CO_2_ and 95% air.

### Viral production

For AAV production, HEK293 cells were transfected with vector plasmids including pAAV encoding an opsin, pHelper and pAAV-RC (serotype 9 or DJ), using a standard calcium phosphate method. After three days, transfected cells were collected and suspended in lysis buffer (150 mM NaCl, 20 mM Tris pH 8.0). After four freeze-thaw cycles, the cell lysate was treated with 250 U/ml benzonase nuclease (Merck) and 1 mM MgCl_2_ for 10−15 min at 37 °C and centrifuged at 4000 rpm for 20 min at 4°C. AAV was then purified from the supernatant by iodixanol gradient ultracentrifugation. The purified AAV solution was concentrated in PBS via filtration and stored at −80°C.

### GloSensor assay

HEK293 cells were cultured in Eagle’s minimal essential medium containing L-glutamine and phenol red (Wako) supplemented with 10% (vol/vol) FBS and penicillin-streptomycin. The cells were co-transfected with the BeGC1 plasmid and the pGloSensor-42F cGMP vector (Promega) using Lipofectamine 2000 (Thermo Fischer Scientific). After transfection, 0.5 μM all-*trans*-retinal (Toronto Research Chemicals) was added to the culture medium. Before measurements, the culture medium was replaced with a CO_2_-independent medium containing 10% (vol/vol) FBS and 2% (vol/vol) GloSensor cGMP stock solution (Promega). The cells were then incubated for 2 h at 27°C in the dark. Intracellular cGMP levels were measured by monitoring luminescence intensity using a microplate reader (Corona Electric) at 27°C^41^.

### Stereotactic surgery

AAV-Ef1a-DIO-ChRmine-eYFP (titer: 1.6 × 10^13^ copies/ml), AAV-hSyn1-SIO-stGtACR1-FusionRed (titer: 1.1 × 10^13^ copies/ml) or AAV-CMV-DIO-hrGFP (titer: 1.5 × 10^13^ copies/ml) was injected bilaterally into the VTA (AP: from −3.0 to −3.3 mm, LM: ±0.5 mm, depth: 4.2 mm) of DAT-IRES-Cre mice under ∼1.2% isoflurane anesthesia. Injection volume was 200 nl per site. The mice were kept in their home cages for at least 3 weeks after AAV injection, prior to behavioral or electrophysiological experiments. For injection of SMPs, SMPs (50 mg/ml) were dispersed with Ringer’s solution. The SMPs were bilaterally injected into the VTA with a volume of 600 nl per site at the same coordinates as AAV injections. In some mice, the SMPs were bilaterally injected into total four sites at the VTA (AP: −3.0 mm, LM: ±0.5 and ±0.2 mm, depth: 4.2 mm). An optical fiber (0.4 mm diameter) attached to a stainless steel ferrule (CFM14L10, Thorlabs) was implanted over the VTA using a cannula holder (XCF, Thorlabs). The optical fiber cannula was permanently cemented to the skull.

### Acute slice preparation

Mice were perfused under isoflurane anesthesia with ice-cold dissection buffer containing (in mM): 87 NaCl, 25 NaHCO3, 25 D-glucose, 2.5 KCl, 1.25 NaH_2_PO_4_, 0.5 CaCl_2_, 7 MgCl_2_, and 75 sucrose, aerated with 95% O2 + 5% CO_2_. The mice were then decapitated, and the brain was isolated and cut into 200-μm-thick horizontal sections on a vibratome in the ice-cold dissection buffer. The slices containing the VTA were incubated for 30 min at 35°C in the dissection buffer and maintained thereafter at RT in standard artificial cerebrospinal fluid (aCSF) containing (in mM): 125 NaCl, 25 NaHCO_3_, 25 D-glucose, 2.5 KCl, 1.25 NaH_2_PO_4_, 1 MgCl_2_, and 2 CaCl_2_), aerated with 95% O2 and 5% CO_2_.

### *In vitro* electrophysiology

Whole-cell patch-clamp recordings from cultured HEK 293 cells or VTA-DA neurons in acute brain slices were performed using an IPA amplifier (Sutter Instruments) at RT. Fluorescently labelled cells were visually identified using an upright microscope (BX51WI; Olympus) equipped with a scientific complementary metal-oxide-semiconductor (sCMOS) video camera (Zyla4.2plus; Andor). The recording pipettes (5−7 MΩ) were filled with the intracellular solution containing (in mM): 135 potassium gluconate, 4 KCl, 4 Mg-ATP, 10 Na2-phosphocreatine, 0.3 Na-GTP, and 10 HEPES (pH 7.3, 280 mOsmol/l). Patch pipettes (5–7 MΩ) had a series resistance of 6.5–25 MΩ, which was compensated to have a final value of 6.5–7.0 MΩ for voltage-clamp recordings^42^. For measuring photocurrents of depolarizing opsins, HEK 293 cells or DA neurons were voltage-clamped at −60 mV. For measuring the photocurrents of GtACR1 and GtACR2, cells were voltage-clamped at 0 mV (HEK 293 cells) or −30 mV (DA neurons). For measuring the photocurrents of ArchT or eNpHR3.0, cells were voltage-clamped at −20–0 mV to achieve near zero holding currents. For current-clamp recordings from DA neurons, the membrane potentials were held at around −60 mV to prevent spontaneous firing unless otherwise noted. For testing the inhibitory effect of stGtACR1 activation (Fig. 2, h and i), spikes at 5–10 Hz were evoked by current injections. In some experiments (Fig. 2, e, f, j and k, and Supplementary Fig. 5), DA neurons were current-clamped at around −40 mV to induce spontaneous APs. In such a case, some DA neurons were highly adaptive to the depolarized membrane potentials and showed little spontaneous firings. We therefore chose the cells that robustly exhibited spontaneous AP rate of more than 1 Hz for further analyses. The PL intensity of the specimen was measured using a photodiode sensor that was routinely calibrated for each experiment.

### Conditioned place preference test

More than 2 weeks after AAV injection, mice were bilaterally injected with SMPs (50 mg/ml in Ringer’s solution) and used for behavioral tests at >1 week after SMP-injection. Conditioned place preference (CPP) tests were performed in an X-ray machine (MX-160Labo, mediXtec Japan) at the dark period. We used two types of test chambers. Both chambers had two compartments with different floor textures, only one of which was irradiated with X-rays. To restrict the X-ray irradiation (X-irradiation) to one side, the other compartment was shielded with lead boards (except for a small gate between the two compartments). Because phasic stimulation to VTA-DA neurons is known to be effective to induce place preference^25^, we designed one of the chambers to introduce pulsed X-irradiation through an X-ray chopper wheel in one component (Fig. 5b: “Chamber I”). We designed another chamber to introduce continuous X-irradiation in one compartment (Fig. 5f) because tonic inhibition of VTA-DA neurons effectively induces place aversion^27^. The test chamber for continuous X-irradiation had different visual cues on the wall of each compartment in addition to different floor textures (Fig. 5f: “Chamber II”).

For CPP tests using ChRmine-expressing mice and corresponding control mice, mice were first habituated to the Chamber I (15 min/day, three sessions). On the first day of the tests, mice were placed in the test chamber and allowed to freely explore for 10 min without X-irradiation. On the following 4 days, the mice were locked in either of the two compartments for 15 min each and received pulsed X-irradiation only in one compartment (150 kV, 3 mV; 50 ms duration,10 pulses at 10 Hz, every 30 s), which we call “Pulsed” conditioning or “P.C.” (Fig. 5c). On the sixth day, the mice were placed in the Chamber I and freely explore for 10 min without X-irradiation. For CPP tests using stGtACR1-expressing mice and corresponding control mice, the mice were habituated to the Chamber II for 5 min and then allowed to explore freely for 10 min without X-irradiation at the first day. On the second and third days, the mice were conditioned for 10 min with freely moving in the chamber where only one of the compartments received continuous X-irradiation (150 kV, 3 mA), which we call “Free moving” conditioning or “F.C.” (Fig. 5f). On the fourth day, the mice were placed in the Chamber II for 10 min without X-irradiation. On the first and last day of the tests, the mice were videotaped at an oblique angle using a USB camera while the whole chamber was illuminated by ambient white LED light, and the ears of mice were tracked offline using DeepLabCut^43^. After completion of the CPP tests, the mice were perfused with 4% PFA and the brain was post-fixed overnight. The brain was cut into coronal sections (section thickness: 80 μm) and the SMP injection sites were observed. The traces of the SMP injections (diameter: 158 ± 10 μm, *n* = 4) were found at ±0−200 μm away from the dorsal edge of VTA. In some cases, these injected SMPs were found along the injection track as well (Supplementary Fig. 9a). Two mice which showed a biased preference for one chamber (>85 %) at the first day were excluded from analysis.

### Immunostaining

For immunostaining of brain slices, we performed transcardial perfusion and post-fixation for overnight using 4% paraformaldehyde (PFA). The fixed brains were sectioned into coronal slices on a vibratome (section thickness: 80 μm) or using a cryostat (after immersion of the fixed brain in 30% sucrose solution for >2 days at 4°C; section thickness: 40 μm). The slices were washed three times with a blocking buffer containing 1 % bovine serum albumin (BSA) and 0.25 % Triton-X in phosphate buffer saline (PBS), and then incubated with primary antibodies (anti-tyrosine hydroxylase, rabbit polyclonal, 1:1000, Merck Millipore; anti-Iba1, rabbit monoclonal, 1:500, Wako; anti-GFAP, mouse monoclonal, 1:1000, Merck Millipore; anti-NeuN, mouse monoclonal, 1:500, Merck Millipore; anti-mouse serum albumin, goat polyclonal, 1:1000, Abcam; anti-doublecortin, rabbit polyclonal, 1:1000, Abcam; anti-c-Fos, rabbit monoclonal, 1:1000, Abcam) in the blocking buffer overnight at 4°C. Only for immunostaining of NeuN, the slices were incubated with the primary antibody for ∼35−40 h at 4°C. The slices were then washed three times with the blocking buffer and then incubated with secondary antibodies (CF594- or CF488A-conjugated donkey anti-rabbit IgG, 1:1000, Biotium; CF488A-conjugated donkey anti-mouse IgG, 1:1000, Biotium; CF488A-conjugated donkey anti-goat IgG, 1:1000, Biotium) in the blocking buffer for 1–2 h at RT. Cellular nuclei were stained by incubation for 10– 15 min with DAPI (2 μM in phosphate buffer) or Hoechst 33342 (5 μg/ml in PBS) at RT. The stained samples were mounted using DABCO, and observed under a florescence microscope (BZ-9000, Keyence) or confocal microscope (LSM710, Zeiss).

### *In vivo* X-irradiation

For cFos induction experiments, more than 2 weeks after AAV injection in the VTA of DAT-IRES-Cre mice, SMPs (50 mg/ml in Ringer’s solution) were injected at the same location as AAV injection. More than 1 week after SMP-injection, mice were anesthetized with a combination anesthetic (0.3 mg/kg medetomidine hydrochloride, 4 mg/kg midazolam, 5 mg/kg butorphanol tartrate) and placed on a heating pad in the X-ray machine. The head of the mice was targeted for X-irradiation at the dose rate of 0.5 or 1 Gy/min (150 kV, 3 mA; 1 min pulses, every 2 min, 5 times). Mice were perfused with 4% PFA at 2 h after X-irradiation.

For assessment of radiation toxicity, mice were irradiated with X-rays following three protocols as described below: (1) Pulsed X-ray radiation: mice were placed in the X-irradiated compartment of Chamber I and received pulsed X-irradiation (150 kV, 3 mA, ∼0.5 Gy/min; 50 ms, 10 Hz, 10 pulses/train, every 30 s; 30 trains/day, 4 days), corresponding to the “Pulsed” conditioning; (2) Fractionated X-ray radiation: mice were placed in the X-irradiated compartment of Chamber II and received fractionated X-irradiation corresponding to the “Free moving” conditioning (150 kV, 3 mA, ∼0.7 Gy/min; 1 min pulses, every 2 min; 5 pulses/day, 2 days); and (3) Acute high-dose radiation: mice were placed in a small box to receive a high dose X-irradiation (150 kV, 3 mA, 1.35 Gy/min; 400 s pulse, once). Total radiation dose was estimated as ∼0.5 Gy for pulsed X-ray radiation, ∼7 Gy for fractionated X-ray radiation and 9 Gy for acute high-dose radiation. Control mice did not receive any radiation.

### EdU staining

To identify newly generated cells in the hippocampus, 5-ethynyl-2’-deoxyuridine (EdU) staining was performed using Click-iT EdU Alexa Fluor 488 Imaging Kit (Thermo Fisher Scientific) following the manufacturer’s protocol. EdU dissolved in Ringer’s solution (10 mg/ml) was intraperitoneally administered for 6 consecutive days starting at 28 days after the last fraction of X-irradiation. The mice were transcardially perfused with 4 % PFA at 28 days after the first day of EdU injection. The brain was post-fixed for overnight and then immersed in 30% sucrose solution for >2 days at 4°C. Coronal sections (thickness: 40 μm) were cut using a cryostat. The sections were incubated with 0.5% TritonX-100 in PBS for 30 min at RT and washed 2 times with PBS containing 3% BSA. The sections were then incubated with Click-iT reaction cocktail for 60 min at RT and washed 3 times with PBS containing 3% BSA. The sections were subsequently subjected to immunostaining of Iba-1 and NeuN with the protocol described above.

### TUNEL assay

For detection of apoptotic cells, TdT-mediated dUTP Nick-End Labeling (TUNEL) assay was performed using Click-iT Plus TUNEL Assay with Alexa Fluor 488 (Thermo Fisher Scientific) following the manufacturer’s protocol. Briefly, mice were transcardially perfused with 4 % PFA and the brain was post-fixed for overnight. The brain was then immersed in 30% sucrose solution for >2 days at 4°C. The brain was cut into coronal sections using a cryostat (section thickness: 10 μm). The sections were permeabilized with proteinase K for 20 min at 37 °C and then incubated with the TdT reaction buffer for 20 min at 37 °C. The sections were then incubated with the TdT reaction mixtures for 120 min at 37 °C, followed by blocking with 3 % BSA and 0.5 % Triton X-100 in PBS for 20 min at RT. The section was then incubated with the Click-iT Plus TUNEL reaction cocktail for 60 min at 37 °C. For DNA staining, the sections were incubated with a Hoechst 33342 solution (5 μg/ml in PBS) for 15 min at RT. After washing, the tissue sections were mounted using DABCO.

### FACS analysis of bone marrow cells

Two long bones (femur and tibia) were isolated per mouse and bone marrow cells were flushed out with 10 ml of FACS buffer (2 % FBS, 2 mM EDTA in PBS) using a syringe with a 25G needle. Cells were treated with 1 ml of ACK buffer (150 mM NH4Cl, 10 mM KHCO3, 0.1 mM Na2EDTA in water) for 1 min, centrifuged at 1500 rpm for 3 min and resuspended in 2 ml of FACS buffer. Live cells were counted with 0.4% (w/v) Trypan Blue Solution (FUJIFILM Wako Chemicals) and were stained at 20 μl of FACS buffer per 1×10^6^ cells containing the first set of following antibodies (all from Biolegend, unless specified otherwise): biotinylated hematopoietic lineage antibodies against NK1.1 (clone PK136), CD11b (M1/70), Ter119 (Ter119), Gr-1 (RB6-8C5), CD4 (GK1.5), CD8α (53-6.7), CD3ε (145-2C11), B220 (RA3-6B2), and IL-7Rα (SB/199) (all 1:500 dilution); BV510-conjugated anti-CD16/32 (93, 1:50), PE-Cy5-conjugated anti-CD135 (A2F10, 1:100), and AF700-conjugated anti-CD48 (HM48-1, 1:100) in FACS buffer for 30 min on ice. Cells were washed with FACS buffer once and stained with the second set of following antibodies: Pacific blue-conjugated streptavidin (1:100, Thermo Fisher Scientific), AF488-conjugated anti-CD150 (TC15-12F12.2, 1:100), PE-conjugated anti-EPCR (eBio1560, 1:100, Thermo Fisher Scientific), PE-Cy7-conjugated anti-c-Kit (2B8, 1:100), APC-conjugated anti-CD34 (HM34, 1:50) in FACS buffer for 90 min on ice. Tubes were tapped every 30 minutes to ensure dispersion of antibodies. Cells were washed twice and resuspended in FACS buffer containing Hoechst (2 μg/ml, Thermo Fisher Scientific) and filtered with a 77μm nylon filter prior to acquisition with the flow cytometer FACSAria™ III (Becton, Dickson and Company). Data were analyzed using the FlowJo software (Becton, Dickson and Company).

### Data analysis

The number of HEK 293 cells in 35-mm culture dishes (Supplementary Fig. 8) was counted by randomly selecting 100 μm × 100 μm squares surrounding the Ce:GAGG crystal for the dishes containing the crystal or anywhere on the coverslip for control dishes. The mean cell density was calculated as the average of the cell densities of three sites per dish.

To estimate the survival rate of dissociated hippocampal neurons (Fig. 4a), we randomly selected three 0.2 mm × 1.0 mm rectangles per dish, surrounding the Ce:GAGG crystal (one of the longer sides of the rectangle was attached to the edge of the crystal) for the dishes containing the crystal or anywhere on the coverslip for control dishes. The dissociated neurons within these sites at DIV1 were monitored at DIV2, 4, and 7.

To quantify the glial accumulation in epi-fluorescence images (Supplementary Fig. 9), binary images were obtained by thresholding. A 100 μm × 100 μm square was drawn around the trace of the SMPs or the optical fiber (one of the sides of the square was attached to the edge of the trace; three sites per image), and the number of pixels exceeding the threshold in these squares was calculated. Likewise, the average number of NeuN-positive cells (Fig. 4b,c) in 100 μm × 100 μm squares drawn around the SMP/optical fiber trace in confocal images (three sites per image) was calculated.

To analyze animal movements in the CPP tests using DeepLabCut^43^, the left and right ears of mice were manually annotated using 20−80 frames per movie to train a deep neural network. An estimated ear position with low likelihood (<0.8) was omitted and replaced with a pixel value obtained using linear interpolation of neighboring values. The middle pixel coordinate between the tracked left and right ears was considered the animal trajectory, assuming that the coordinate represented the location of the mouse head. The heat maps shown in Fig. 5e,i indicate the probability of the presence of the animal trajectory within each 100 × 100 pixel image. The number of rearing events was calculated by counting the number of times that the height of the mouse head exceeded a threshold level. In the CPP tests, two mice which showed a substantially biased preference for one of the two compartments (>85 %) at the first day were excluded from the analysis.

All values are expressed as mean ± SEM. Statistical tests were performed using GraphPad Prism or Igor Pro. The normality of data distribution was routinely tested. Analyses of two sample comparison were performed using unpaired or paired *t*-tests when each sample was normally distributed, or Mann-Whitney U tests when at least one of the samples in every two-sample comparison was not normally distributed. Tests for two-sample comparison were two-sided. Statistical analyses for multiple comparisons were carried out using one- or two-way ANOVA followed by Bonferroni’s multiple comparison tests or Dunnett’s multiple comparison tests vs. the control, unless otherwise noted.

## Acknowledgements

We thank H. Kasai for providing DAT-IRES-Cre mice; K. Deisseroth for providing bReaChES and ChRmine plasmids; H. Yawo for providing PsChR and C1V1 plasmids; B. Haider for comments; Y. Miyoshi, S. Tsukamoto, A. Kambara, C. Koike, M. Jin for technical assistances. This work was supported by JST-PRESTO (JPMJPR168D) to T.Yamashita; KAKENHI grants (16H05927, 17H05744 and 19H03533) to T.Yamashita; Asahi Glass Foundation to T.Yamashita; Sumitomo Foundation to T.Yamashita; and JST-CREST (JPMJCR1656) to A.Y.

## Author contributions

T.Yamashita conceived and designed the project with inputs from T.M. and T.Yanagida. T.M. conducted animal surgery, behavioral experiments, cell culture and histology experiments, and analyzed data. T.M. and T. Yamashita carried out AAV production and characterization of Ce:GAGG crystals. T.Yanagida and N.K. provided Ce:GAGG crystals and performed spectral measurements. T.N. and J.Y. analyzed behavioral data and provided simulation of RL intensity. S.P.T. and H.K. helped with Glosensor experiments. M.S. and H.T. performed FACS analysis of bone marrow cells; S.H. helped with constructing plasmids. S.U. and S.T-K. helped with implantation of Ce:GAGG crystals. A.Y. provided plasmids and AAV and contributed to interpretation of data. T.Yamashita performed all physiological measurements, Glosensor experiments, analyzed data, and wrote the manuscript with inputs from other co-authors.

## Competing interests

T.Yamashita, T. M., T. Yanagida, and N. K. filed a patent for optogenetic use of Ce:GAGG.

---

**Supplementary Fig. 1.**
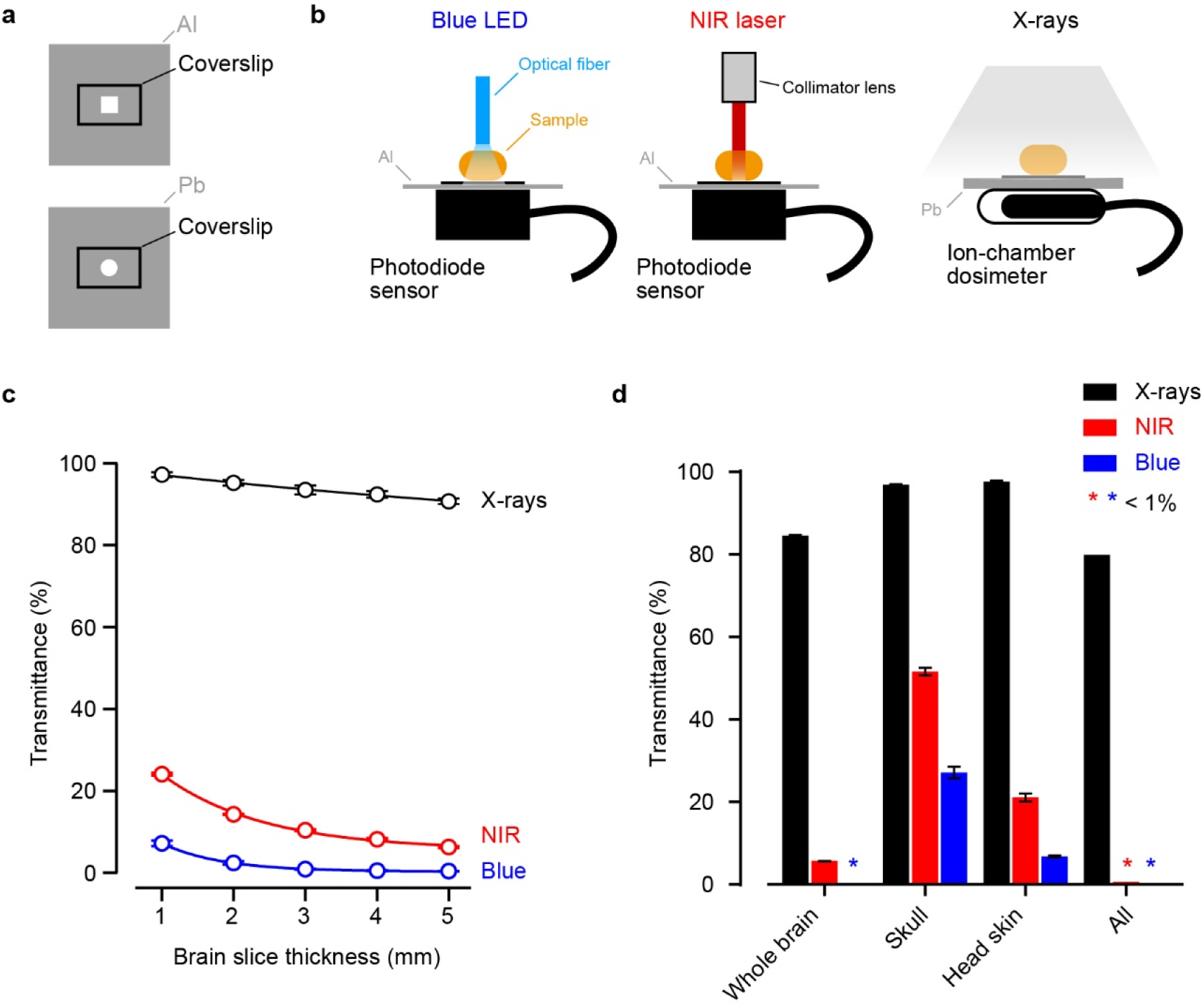
Tissue penetration by X-rays, NIR, and blue light. **a**, The tissue penetration efficacy of different electromagnetic waves was measured using an aluminum (Al, top) or lead (Pb, bottom) plate with a hole shaped as a square (5 mm × 5 mm, top) or a circle (5 mm diameter, bottom) with a coverslip. The lead board was 3 mm thick. **b**, Schematic of the experiments. The photosensor or dosimeter was placed under the hole in the plate. **c**, Transmittance of brain slices of different thickness irradiated with X-rays (black, 150 kV, 3 mA; input: 0.116−0.122 Gy/min), NIR laser (red, 976 nm; input: 42.8−43.9 mW/cm^2^) and blue LED (blue, 470 nm; input: 48.6−49.5 mW/cm^2^) (*n* = 3 slices for each group). **d**, Transmittance of the shaved head skin, skull, and whole brain of mice (*n* = 3 mice for each group). Samples were placed horizontally. Input power: 0.129−0.131 Gy/min for X-rays; 43.1−45.1 mW/cm^2^ for NIR; 51.3 mW/cm^2^ for blue LED. Values are mean ± SEM.

**Supplementary Fig. 2.**
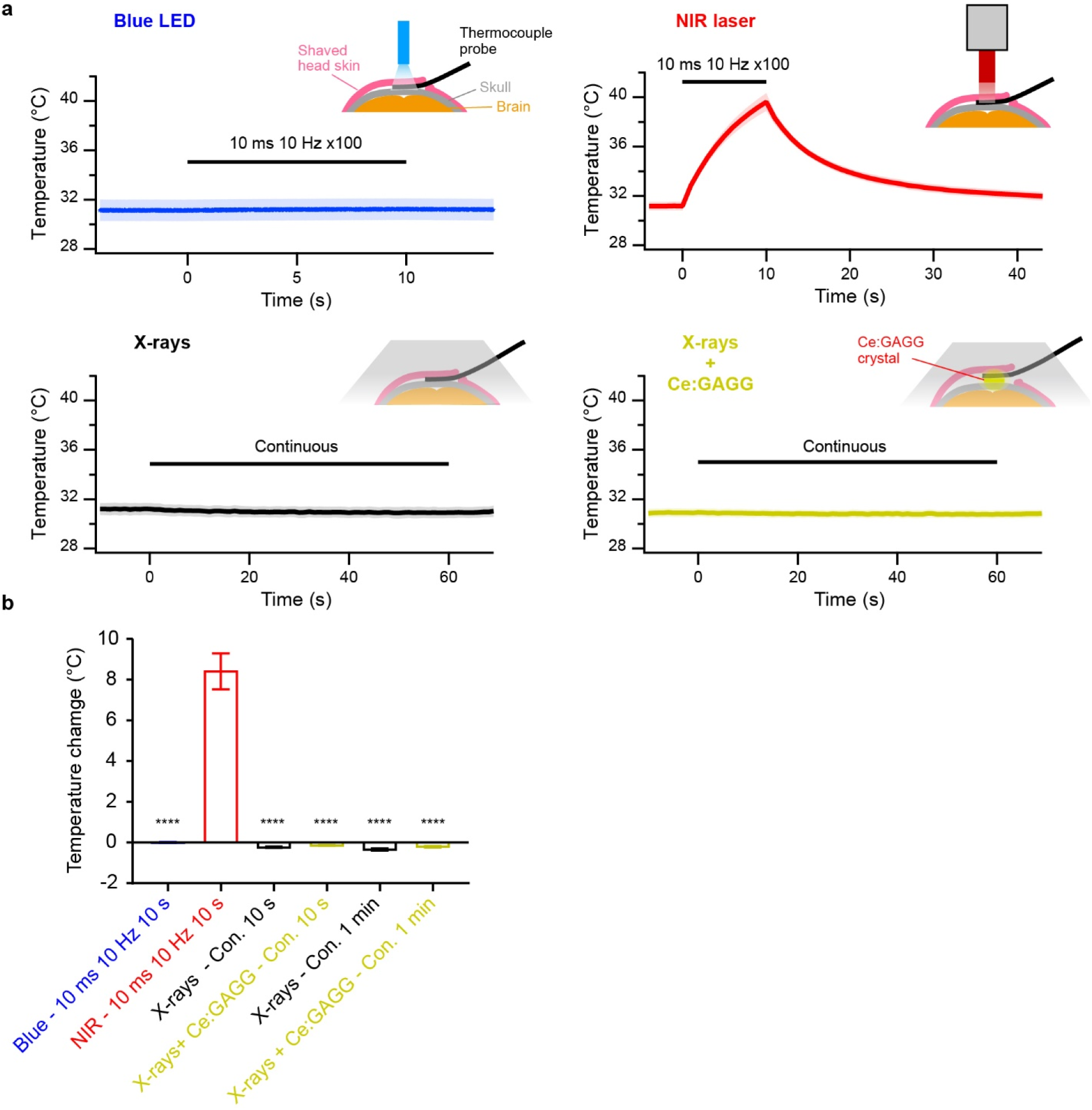
Temperature under the skin of the head irradiated by X-rays, NIR, or blue light. **a**, Average traces of temperature recordings (*n* = 4 mice each). The temperature between the shaved head skin and skull of anesthetized mice irradiated with blue LED (10 mW at the fiber tip; fiber diameter, 400 μm), NIR laser (200 mW/mm^2^ after a collimator lens with a beam diameter of 4 mm) or X-rays (150 kV, 3 mA, 1.35 Gy/min) was measured. Scintillation-induced temperature changes were measured by placing a Ce:GAGG crystal (6 mm × 4 mm × 1 mm) together with the temperature sensor between the skin of the head and the skull. The anesthetized mouse was placed on a heat pad. **b**, Temperature changes under various conditions. Con.: continuous. *F*_5,18_ = 94.7, * * * * *P* < 0.0001, Bonferroni’s multiple comparison test, vs. NIR. Values are mean ± SEM.

**Supplementary Fig. 3.**
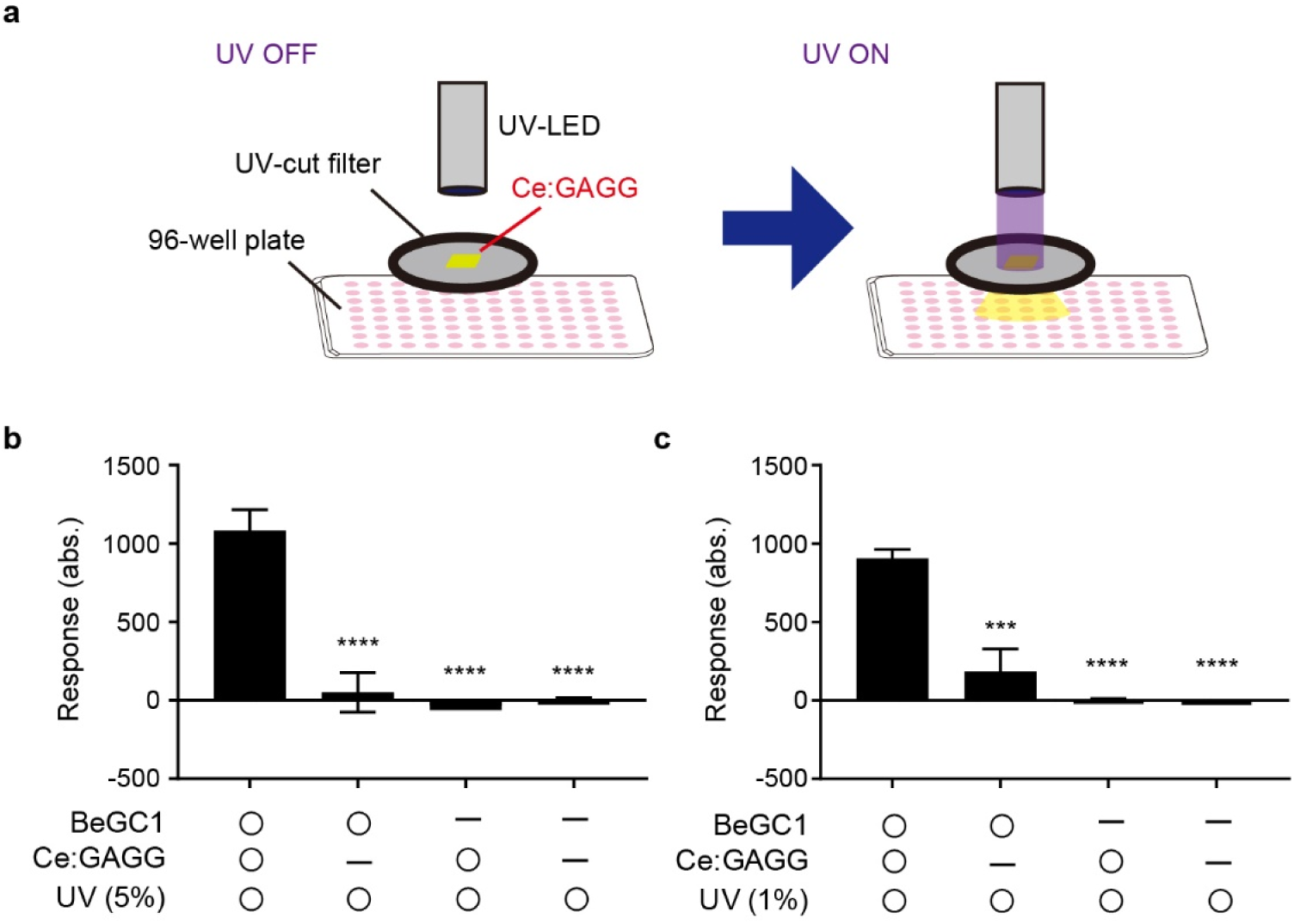
Ce:GAGG photo-luminescence activates the enzyme rhodopsin BeGC1. **a**, Experimental setup. A Ce:GAGG crystal (6 mm × 4 mm × 1 mm) was irradiated by UV (1% or 5% intensity of the maximum LED power) to illuminate HEK 293 cells expressing BeGC1 or control cells cultured in 96-well plates. PL intensity: 1.56 μW/mm^2^ for 5% UV; 0.66 μW/mm^2^ for 1% UV. UV irradiation was largely attenuated by a UV-cut filter placed over the plate. The same wells were irradiated with UV only through the UV-cut filter. **b**, BeGC1 activation was quantified by measuring the luminescence intensity derived from Glosensor (*n* = 4 wells each). Response amplitude was calculated by substituting the intensity at 8−12 min after 5% UV irradiation from the baseline. *F*_3,12_ = 30.9, * * * * *P* < 0.0001, Bonferroni’s multiple comparison test, vs. the BeGC1(+)/Ce:GAGG(+)/UV(+) group. **c**, Same as **b**, but with 1% UV irradiation (*n* = 4 wells each). *F*_3,12_ = 34.2, * * * *P* < 0.001, * * * * *P* < 0.0001, Bonferroni’s multiple comparison test, vs. the BeGC1(+)/Ce:GAGG(+)/UV(+) group. Values are mean ± SEM.

**Supplementary Fig. 4.**
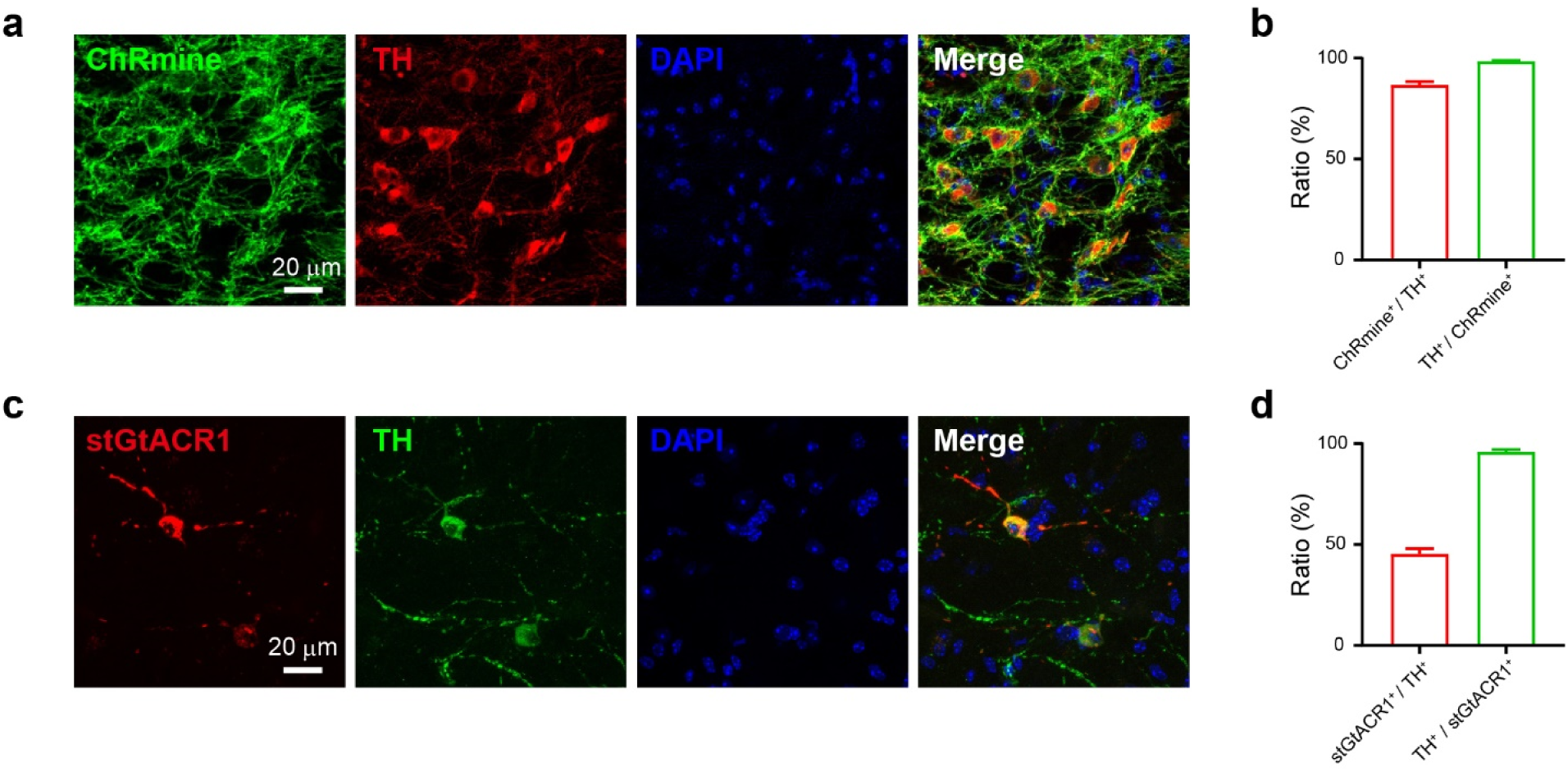
DA neuron-specific expression of ChRmine and stGtACR1. **a**, Representative confocal images showing virally induced expression of ChRmine-eYFP (green) and immunostained TH (red) in the VTA. Blue:DAPI. **b**, Quantification of overlapping of ChRmine-eYFP- and TH-labelled neurons in the VTA (*n* = 3 mice). **c**, Same as **a**, but for stGtACR1-FusionRed (red) and immunostained TH (green). **d**, Same as **b**, but for stGtACR1-FusionRed- and TH-labelled neurons (*n* = 3 mice). Values are mean ± SEM.

**Supplementary Fig. 5.**
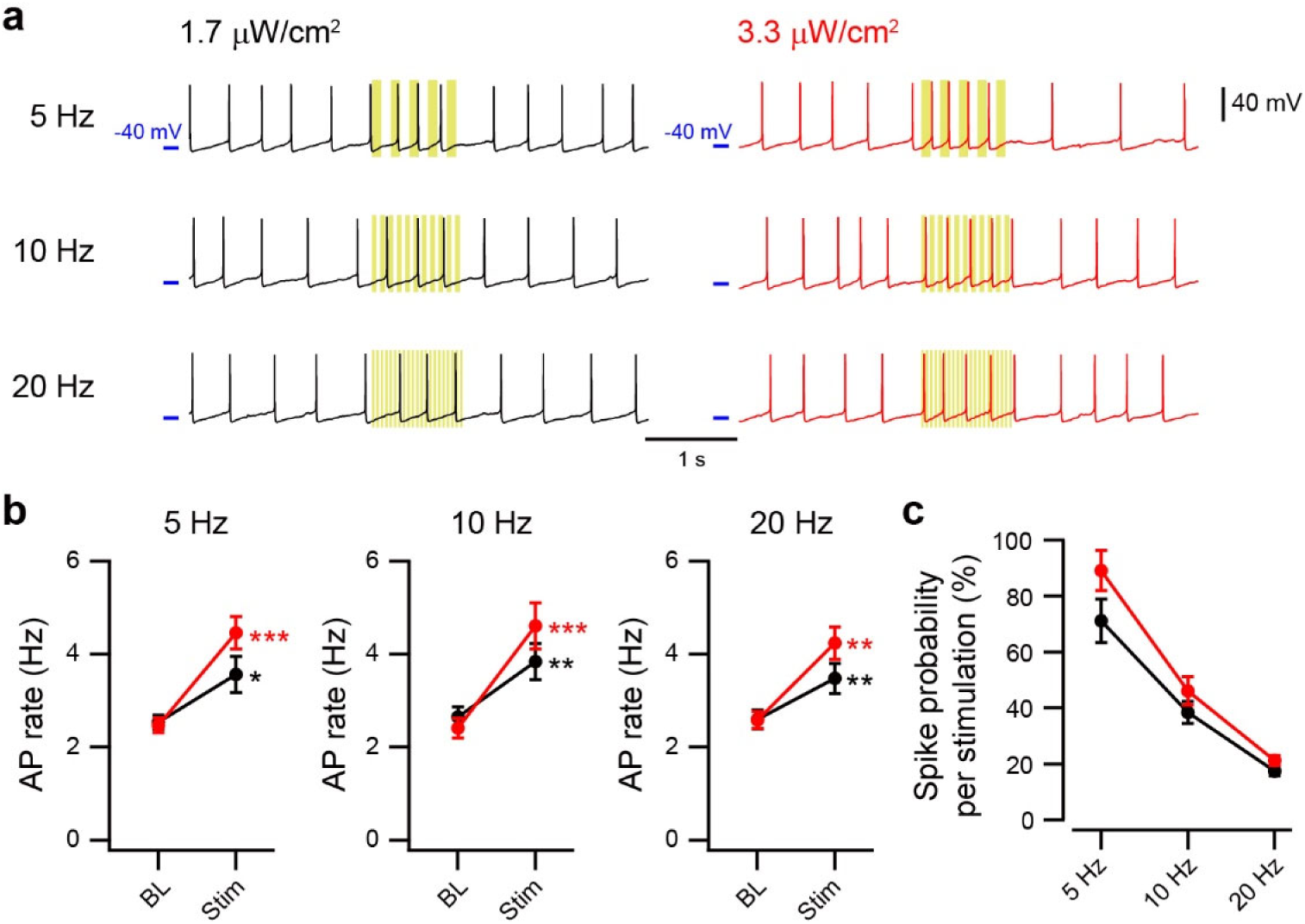
Activation of VTA-DA neurons by low-intensity PL. **a**, Example recordings from a ChRmine-expressing VTA-DA neuron illuminated with pulses of Ce:GAGG PL (yellow bars) at 1.7 (left) and 3.3 (right) μW/cm^2^. 5 Hz: 100 ms, 5 pulses at 5 Hz; 10 Hz: 50 ms, 10 pulses at 10 Hz; 20 Hz: 25 ms 20 pulses at 20 Hz. **b**, Quantification of AP rates at the baseline (BL) and during pulsed PL illumination (Stim, *n* = 7−8 cells) at 1.7 (black) and 3.3 (red) μW/cm^2^. * *P* < 0.05, * * *P* < 0.01, * * * *P* < 0.001, paired *t* tests vs. BL. **c**, Quantification of spike probability upon stimulation at different frequencies (*n* = 7−8 cells). Values are mean ± SEM.

**Supplementary Fig. 6.**
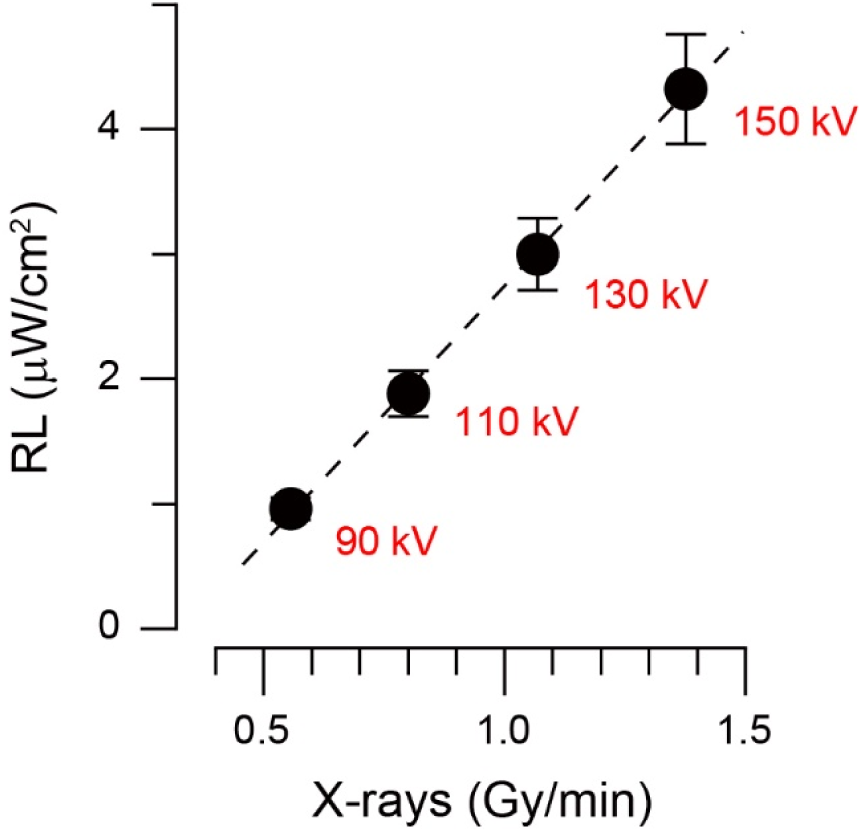
Intensity of Ce:GAGG RL. The intensities of RL emitted by SMPs were measured by fiber optics at different tube potentials of the X-ray source with a constant tube current (3 mA) (*n* = 3 samples). The X-ray dose rates at these tube potentials were measured by a dosimeter. These values in combination with the exponential decay of RL intensities in gray matter (measured as 32.75 % reduction by 200 μm distance) are used for simulation of a 3D map of RL emitted from a spherical aggregate of injected SMPs (Fig. 3b).

**Supplementary Fig. 7.**
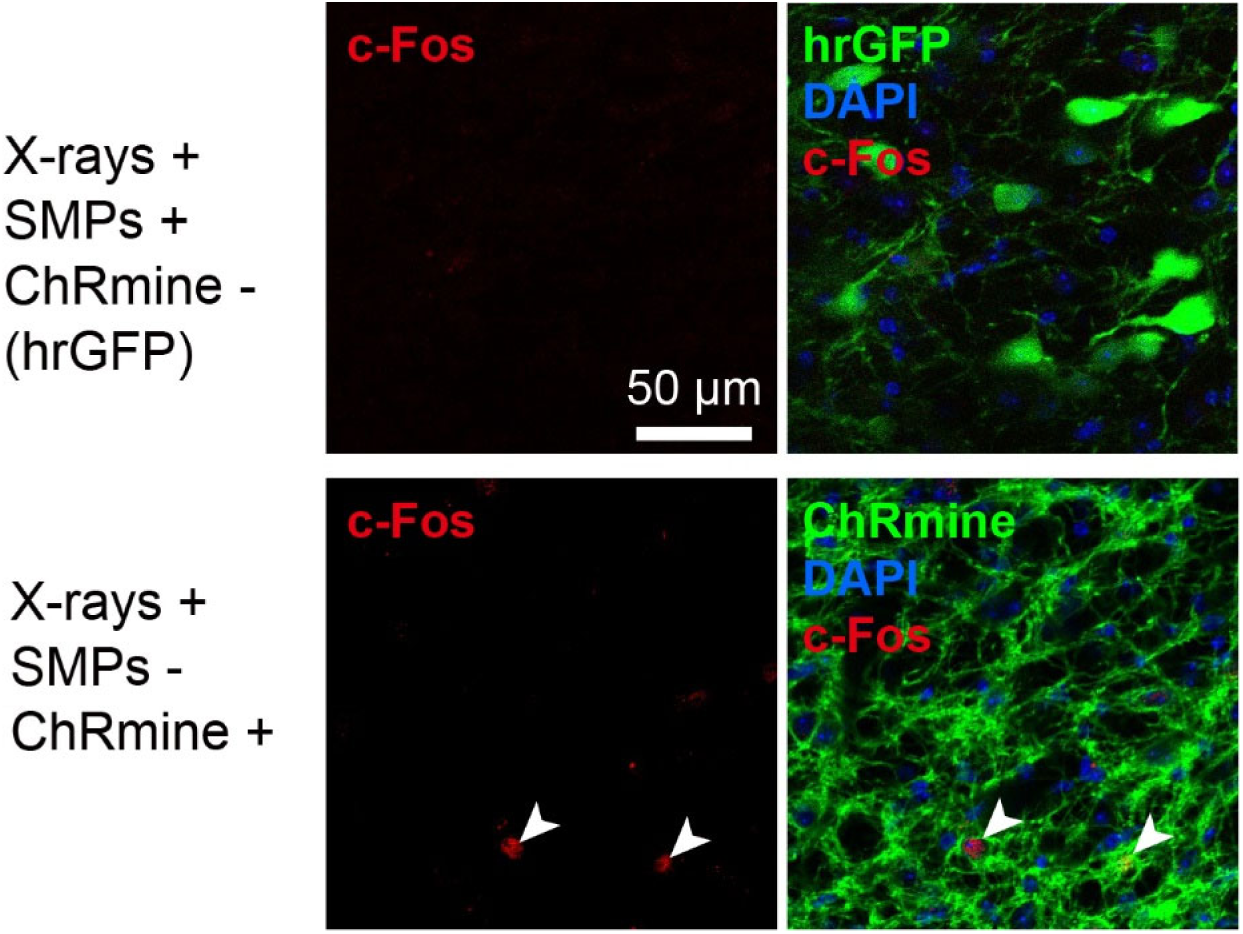
Immunoreactivity against c-Fos in control mice. Confocal images of immunoreactivity against c-Fos (red) of the VTA-DA neurons in control mice irradiated with X-rays (1.0 Gy/min, total 5 min). Representative images from a mouse with SMP injection but no ChRmine expression (top, hrGFP was expressed in the VTA-DA neurons) and that with ChRmine expression but no SMP injection (bottom) are shown.

**Supplementary Fig. 8.**
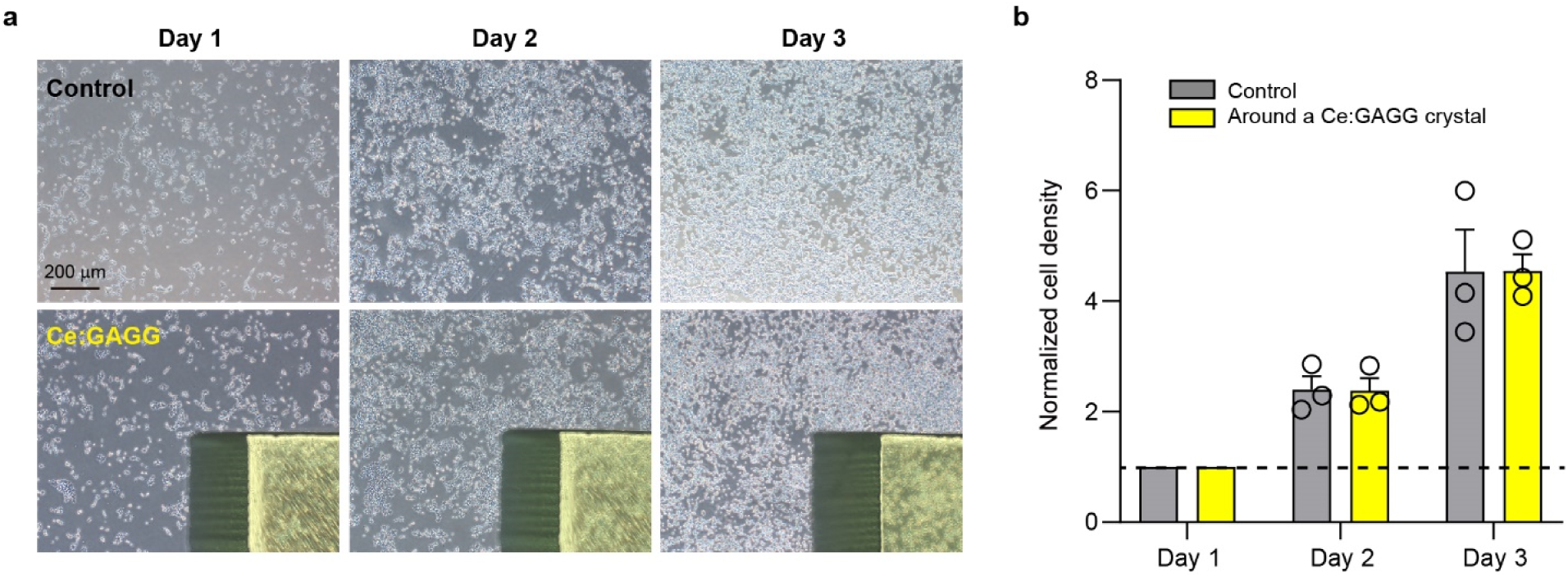
Proliferation of HEK293 cells around a Ce:GAGG crystal. **a**, HEK293 cells were cultured in a dish with (bottom) or without (top) a Ce:GAGG crystal. **b**, During 3 days of culturing, the proliferation rate of the cells at around the Ce:GAGG crystal (*n* = 3 dishes) did not differ significantly from that in the control dish (*n* = 3 dishes, *F*_1,4_ = 0.000023, *P* > 0.99, two-way ANOVA). Values are mean ± SEM.

**Supplementary Fig. 9.**
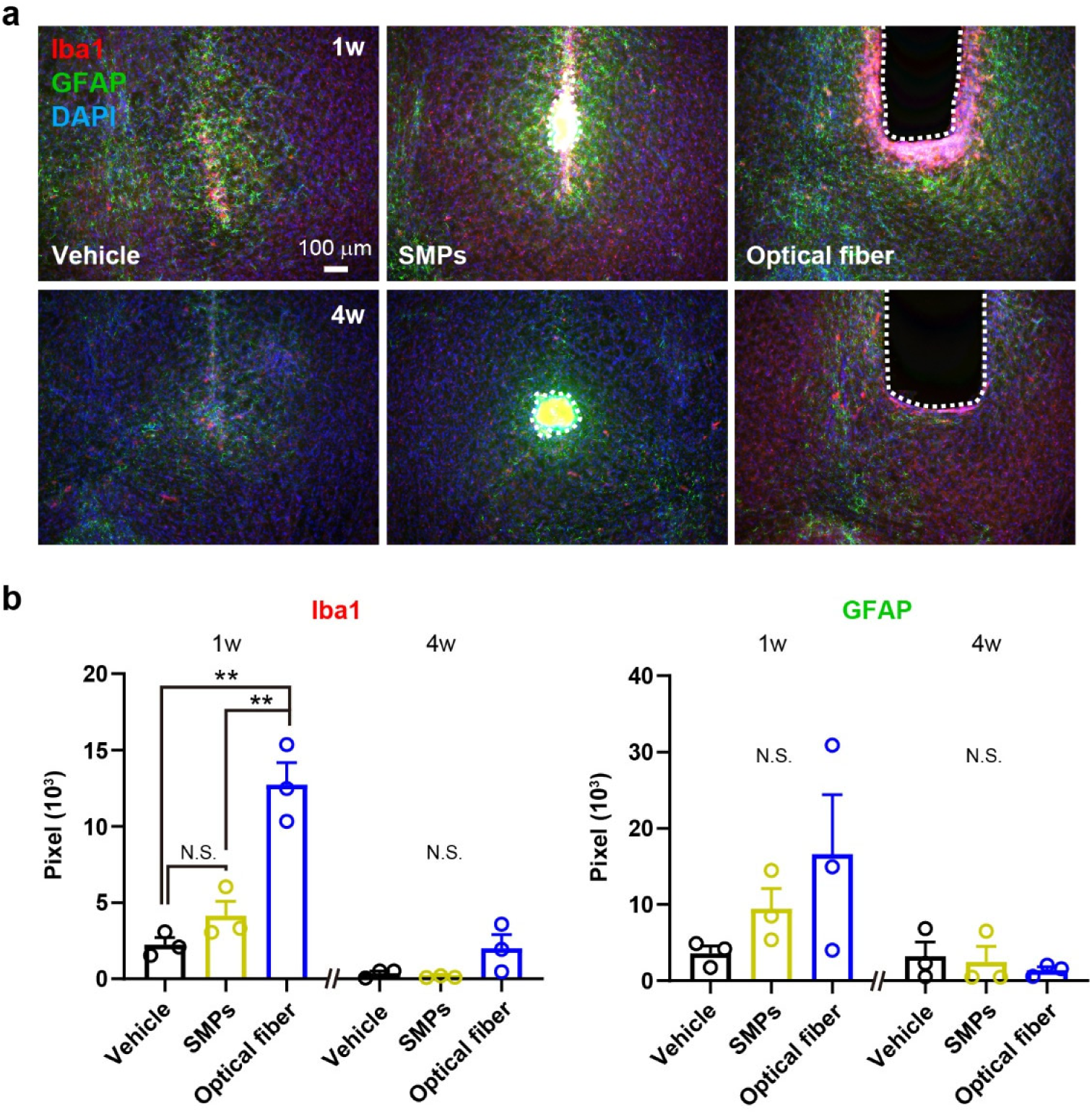
Glial cells around implanted SMPs and optical fibers. **a**, Representative epi-fluorescence images of coronal slices showing immunostaining for activated microglia (Iba1, red) and astrocytes (GFAP, green) at the injection site of vehicle (left) or SMPs (middle), and at the ventral tip of an implanted optical fiber (right). Slices were obtained from the mice at one (1w, top) or four weeks (4w, bottom) after surgery. The trace of SMPs or an optical fiber is outlined by a dashed line. Blue: DAPI. **b**, Quantification of accumulation of microglial cells (Iba1, left) and astrocytes (GFAP, right) estimated in 100 μm x 100 μm squares near the injection/implantation traces (*n* = 3 mice for each group). * * *P* < 0.01, N.S., not significant, Bonferroni’s multiple comparison test for Iba1-1w and one-way ANOVA for Iba1-4w (*F*_2,6_ = 3.76, *P* = 0.088), GFAP-1w (*F*_2,6_ = 1.85, *P* = 0.24), and GFAP-4w (*F*_2,6_ = 0.326, *P* = 0.73). Values are mean ± SEM.

**Supplementary Fig. 10.**
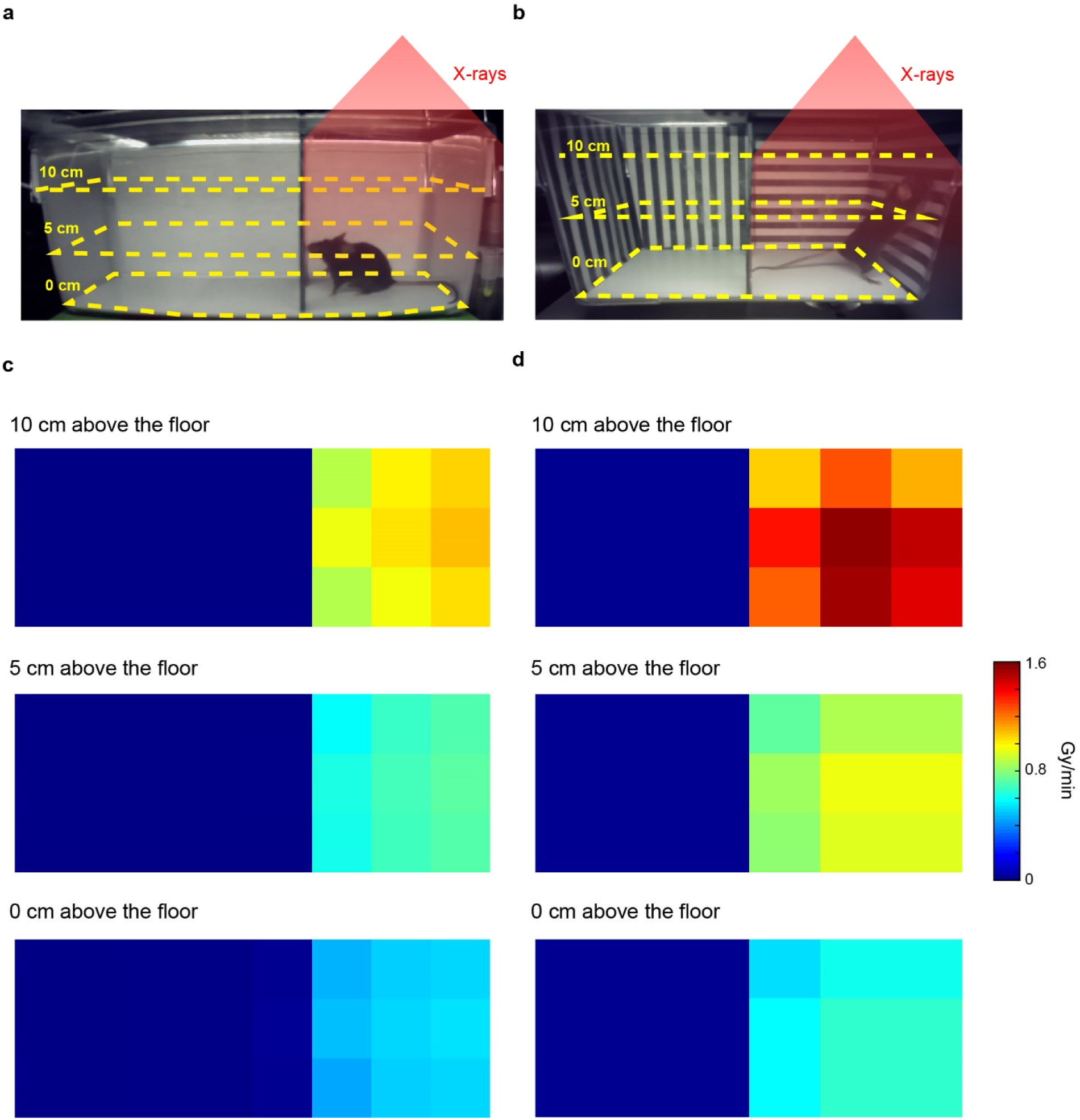
X-ray dose rates measured in the CPP test chambers. **a** and **b**, X-ray dose rates were measured with a dosimeter placed at 0, 5 or 10 cm above the floor of Chamber I (**a**) and Chamber II (**b**). **c**, Heat maps showing X-ray dose rates (in Gy/min) measured in both X-ray irradiated (right) and non-irradiated (left) compartments of Chamber I in the absence of the X-ray chopper. Section size: 3.3 cm × 3.2 cm. **d**, Same as **c**, but for Chamber II. Section size: 3.3 cm × 4.0 cm.

**Supplementary Fig. 11.**
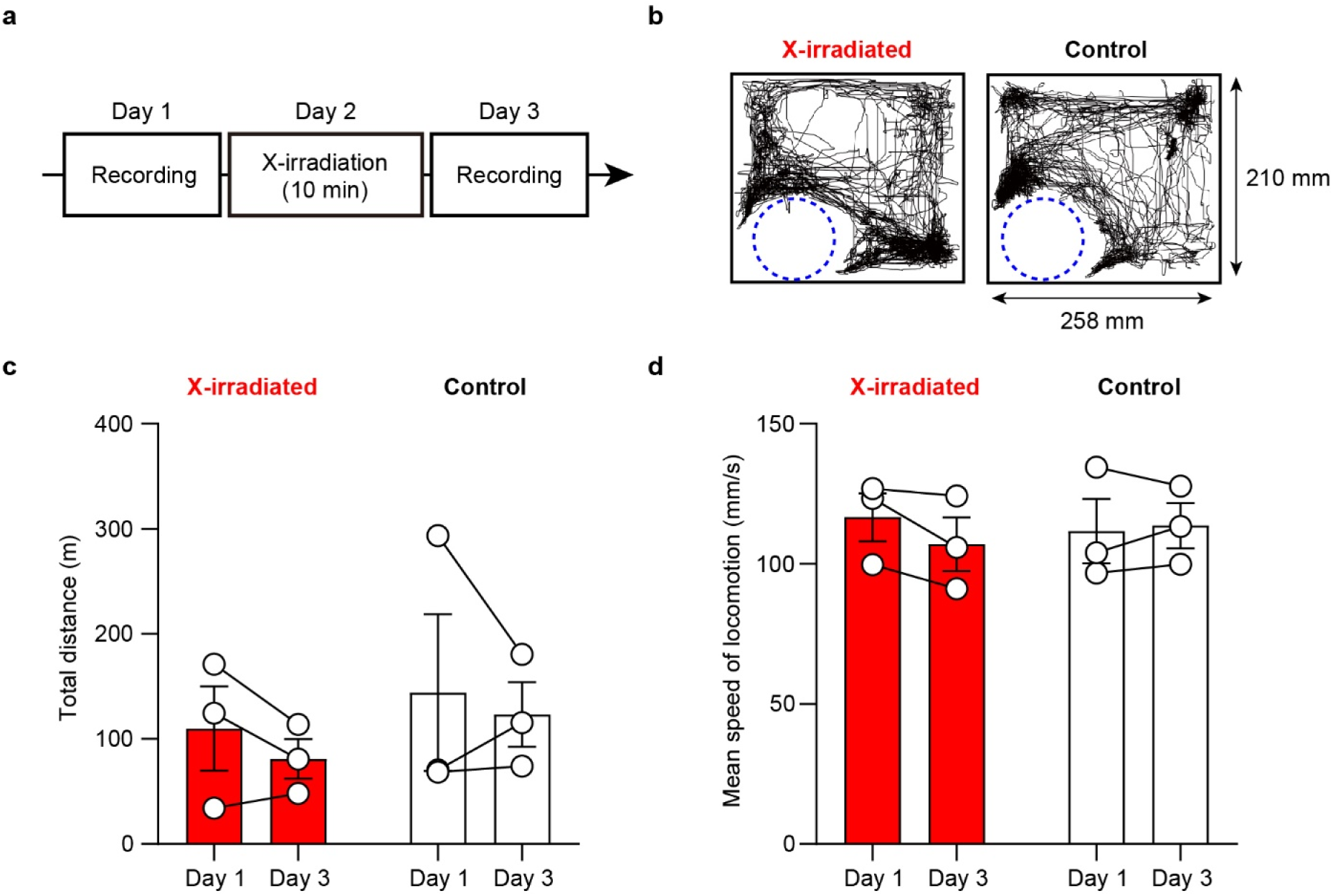
X-irradiation did not affect locomotor behavior at the home cage. **a**, Schematic of the experiment. X-rays (150 kV, 3 mA) was irradiated onto anesthetized mice for 10 min in the X-irradiated compartment of the CPP test chamber for “Free moving” conditioning. **b**, Top-view trajectory of X-irradiated (left) or non-irradiated (right, Control) mice during 1 h recording of their home-cage at day 3. One mouse per cage. The center of the mouse body was tracked using DeepLabCut^43^. The dotted circle indicates the location of a water bottle. **c** and **d**, Total travel distance **c** and mean locomotion speed **d** of the mice at day 1 and day 3 (*n* = 3 mice each). The mean locomotion speed was calculated as the average values in the film frames where the tracked body location moved at >50 mm/s [**c** X-irradiated: *P* > 0.31; Control: *P* > 0.70; **d** X-irradiated: *P* > 0.15; Control: *P* > 0.72; paired *t* test]. Values are mean ± SEM.

**Supplementary Fig. 12.**
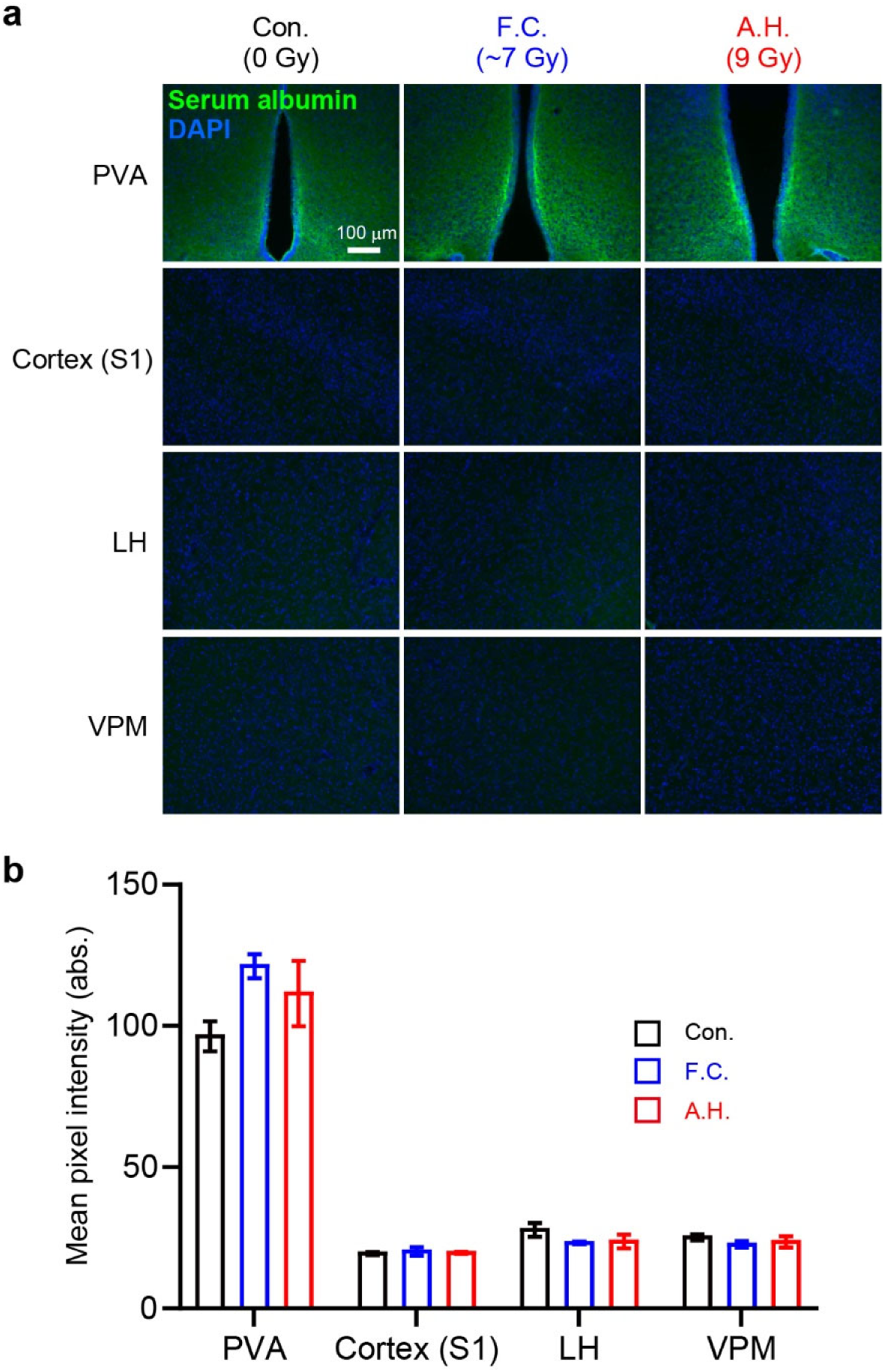
Immunoreactivity against serum albumin after X-irradiation. **a**, To assess the brain blood barrier function, immunoreactivity against extravasated serum albumin was examined in the brain of mice perfused at 3 days after the last fraction of X-irradiation. Representative epi-fluorescence images show immunoreactivity against mouse serum albumin (green) in the paraventricular area (PVA), primary somatosensory cortex (S1), the lateral hypothalamus (LH) and the ventral posteromedial nucleus of the thalamus (VPM), in a mouse that experienced acute high-dose radiation (A.H.), fractionated radiation corresponding to “Free moving” conditioning (F.C.) or no radiation (Con.). **b**, Mean pixel intensities of mouse serum albumin immunoreactivity were not different among three groups in all of the brain areas (PVA, *F*_2,6_ = 2.64, *P* = 0.15; S1, *F*_2,6_ = 0.15, *P* = 0.86; LH, *F*_2,6_ = 1.61, *P* = 0.28; VPM, *F*_2,6_ = 0.85, *P* = 0.47; one-way ANOVA). Values are mean ± SEM.

**Supplementary Fig. 13.**
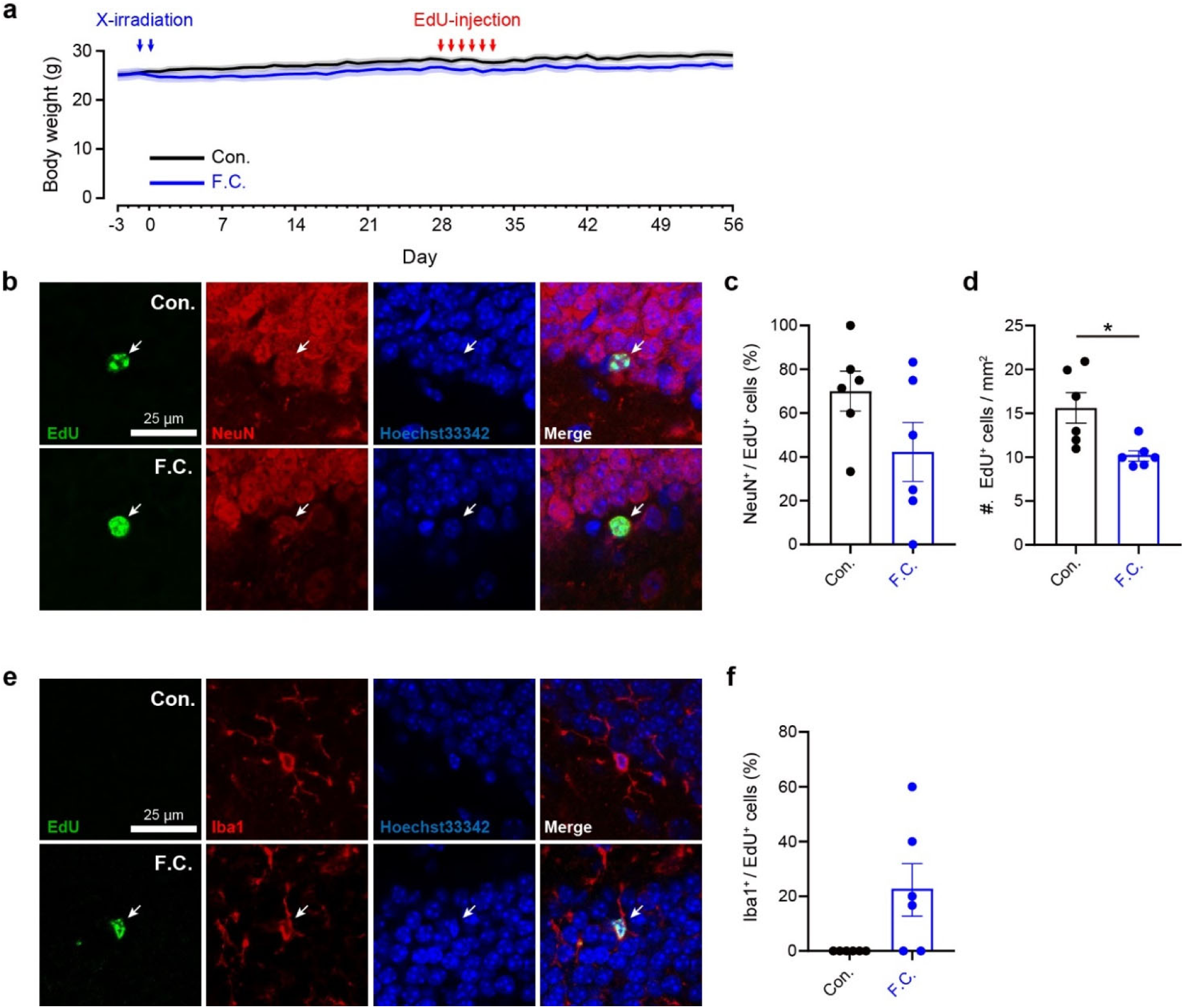
Body weight and neurogenesis in the hippocampal dentate gyrus after X-irradiation. **a**, Body weight of the mice that experienced X-irradiation (blue arrows) with the “Free moving” conditioning protocol (F.C.; blue, *n* = 6 mice) or no radiation (Con.; black, *n* = 6 mice). Red arrows indicate the timings of EdU (10 mg/ml) injection. No significant difference was observed between two groups (*F*_1,10_ = 2.27, *P* = 0.16, two-way ANOVA). All mice survived for 56 days after radiation. **b**, Confocal images showing EdU-positive cells co-expressing NeuN (arrows) at the subgranular zone of the hippocampal dentate gyrus in the X-irradiated (F.C., bottom) or the control (Con., top) mice. The mice were perfused at 28 days after the first day of EdU injection. **c**, Quantification of NeuN-positive cells among EdU-positive cells in the subgranular zone (*n* = 6 mice each, *P* = 0.19, Mann-Whitney U test). **d**, Quantification of EdU-positive cells indicates that the M.C. radiation impaired stem cell proliferation. * *P* < 0.05, Mann-Whitney U test. **e**, Confocal images of Iba1-positive cells at the dentate gyrus of the X-irradiated (F.C., bottom) or non-X-irradiated (Con., top) mice. **f**, In 4 out of 6 X-irradiated mice, Iba1-positive cells were found among EdU-positive cells, whereas no such newly generated microglial cells were found in the control mice. Filled circles indicate individual data. Values are mean ± SEM.

**Supplementary Fig. 14.**
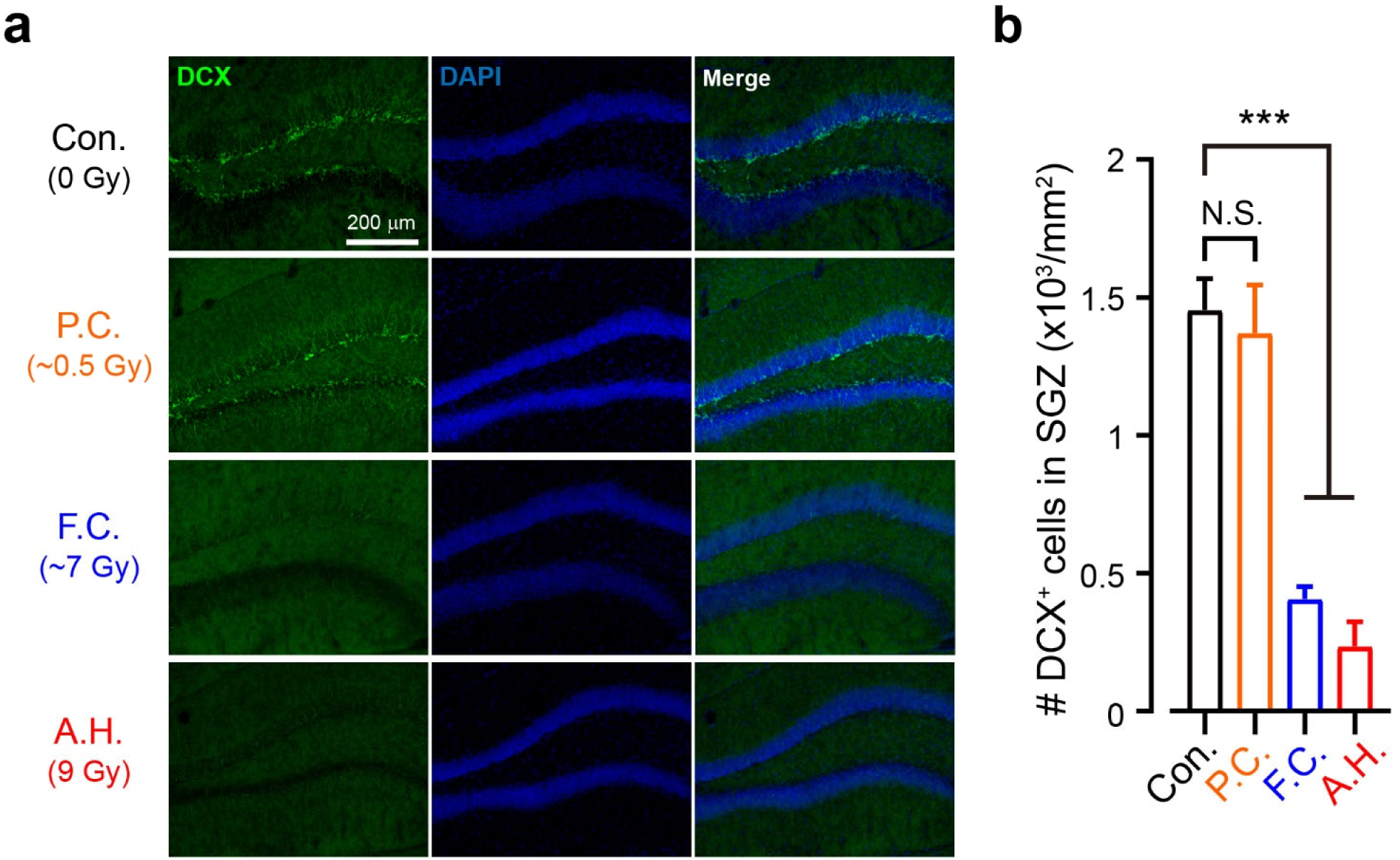
Immature neurons in the hippocampal dentate gyrus after X-irradiation. **a**, Doublecortin (DCX) immunoreactivity (green) at the hippocampal dentate gyrus two days after acute high-dose radiation (A.H.), “Pulsed” conditioning (P.C.), “Free moving” conditioning (F.C.), or no radiation (Con.). Blue: DAPI. **b**, Quantification of DCX-positive cells under different conditions (*n* = 3 mice for each group). * * * *P* < 0.0001, Bonferroni’s multiple comparisons test vs. the control. Values are mean ± SEM. N.S. indicate no significant differences vs. the control.

**Supplementary Fig. 15.**
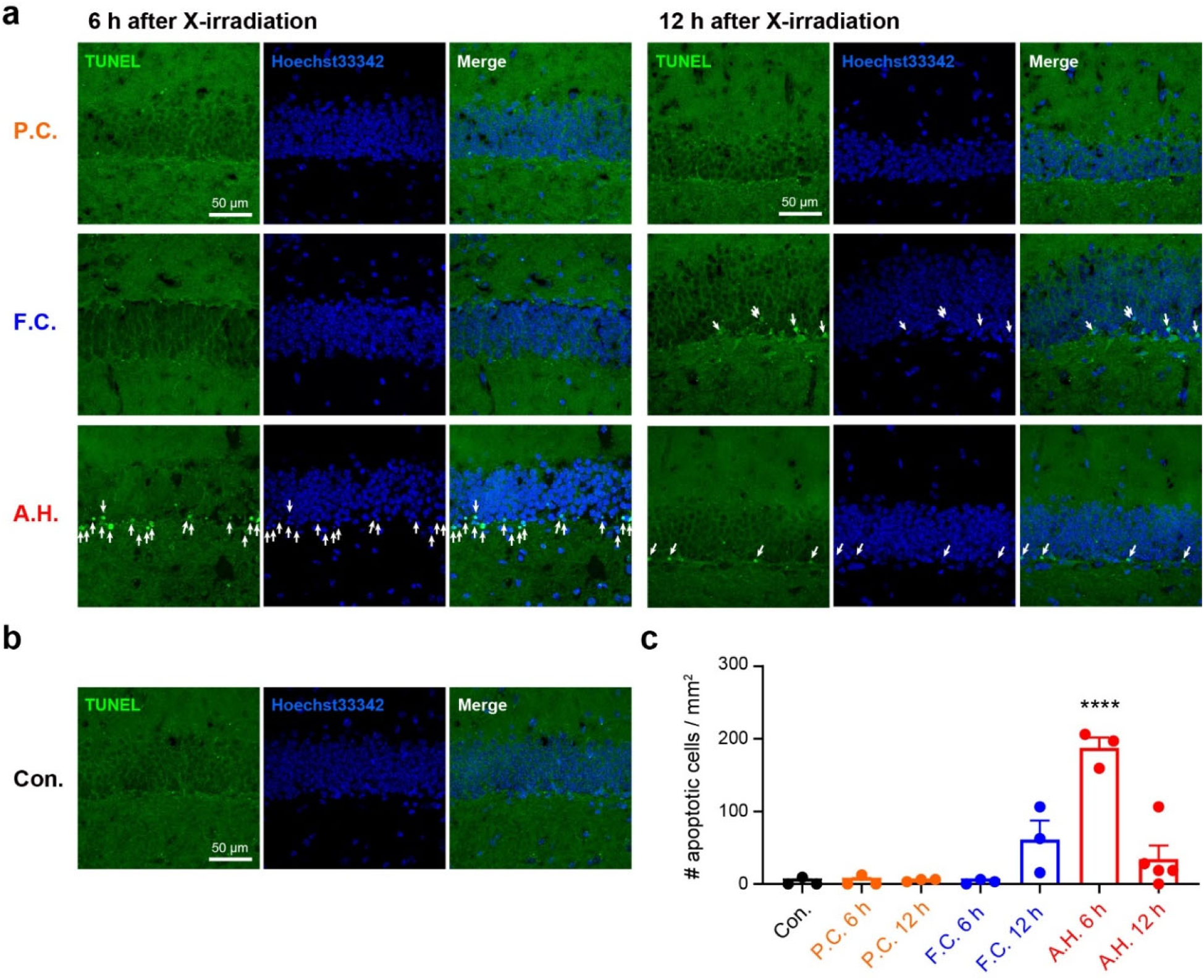
Apoptotic signals in the hippocampal dentate gyrus after X-irradiation. **a**, Confocal images of TUNEL staining in the hippocampal slices obtained from the mice irradiated with X-rays. The mice were perfused 6 (left) or 12 (right) h after the last fraction of X-irradiation. P.C.: “Pulsed” conditioning, F.C.: “Free moving” conditioning, A.H.: Acute high-dose radiation. Arrows indicate apoptotic cells. **b**, TUNEL staining in the non-X-irradiated control (Con.) mice. **c**, Quantification of the number of apoptotic cells in the hippocampal dentate gyrus (*n* = 3-5 mice for each group). Filled circles indicate individual data. * * * * *P* < 0.0001, Bonferroni’s multiple comparisons test vs. the control. Values are mean ± SEM.

**Supplementary Fig. 16.**
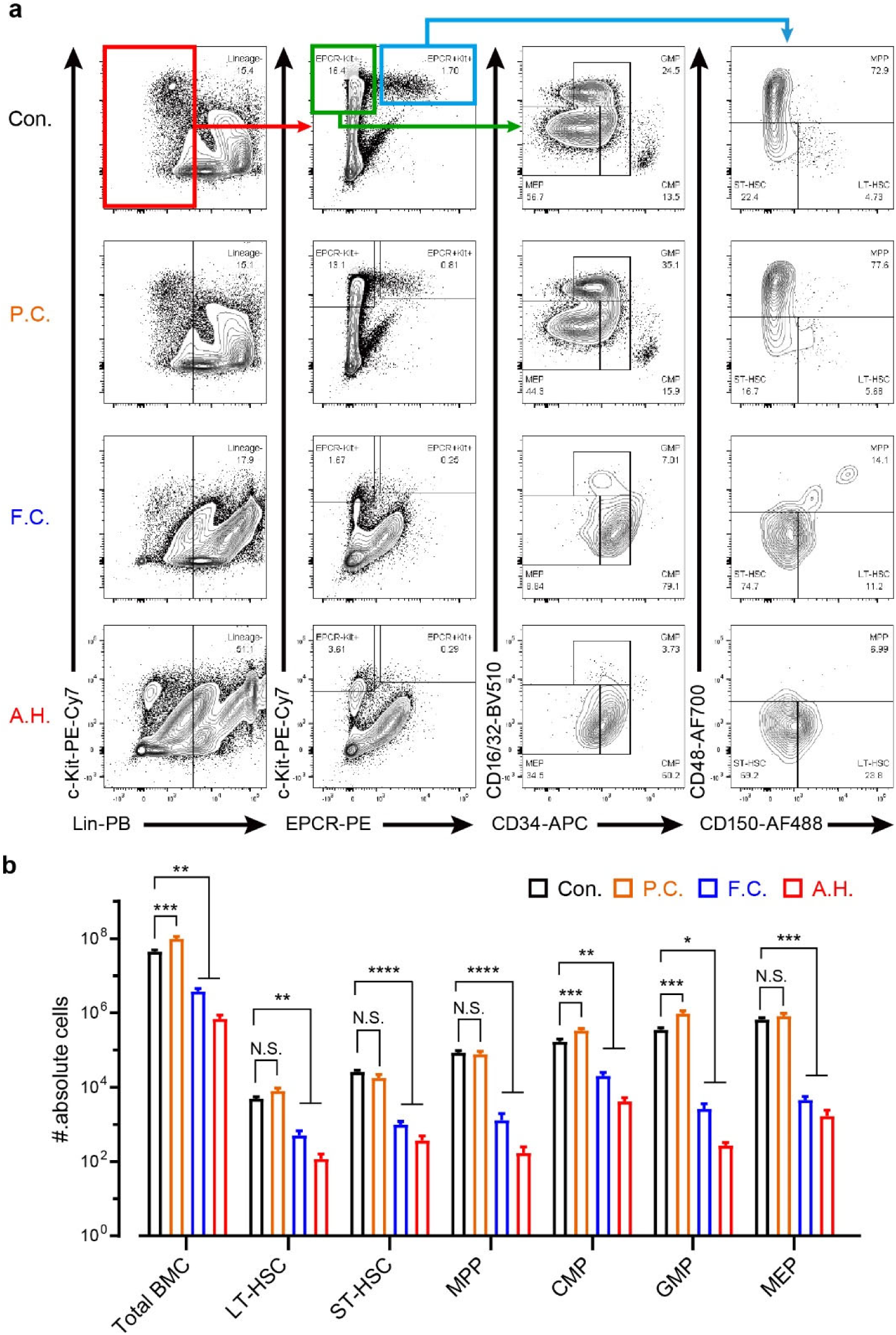
FACS analysis of the bone marrow cells in X-irradiated mice. **a**, Representative FACS plots of the bone marrow obtained at 3 days after the last fraction of X-irradiation. Con.: no radiation control, P.C.: “Pulsed” conditioning, F.C.: “Free moving” conditioning, A.H.: Acute high-dose radiation. **b**, Absolute cell numbers of various cell populations in the bone marrow (*n* = 5 mice for each group). BMC: bone marrow cell, LT-HSC: long-term hematopoietic stem cell, ST-HSC: short-term hematopoietic stem cell, MPP: multipotent progenitor, CMP: common myeloid progenitor, GMP: granulocyte-myeloid progenitor, MEP: megakaryocyte-erythroid progenitor. * *P* < 0.05, * * *P* < 0.01, * * * *P* < 0.001, * * * * *P* < 0.0001, Dunnett’s multiple comparison tests vs. the control. N.S. indicate no significant differences vs. the control.

## Notes

### Summary of Updates

We added data on electrophysiology using ChRmine (Figs. 1 and 2; Supplementary Figs. 4 and 5); we obtained new data using Ce:GAGG microparticles and then replaced and added data related to in vivo validation of our technology (Figs. 3-5; Supplementary Figs. 6, 7, 9, 10 and 12-16); we updated and rewrote the manuscript.

